# Competition between distinct ApoE alleles and mCRP for the endothelial receptor CD31 differentially regulates neurovascular inflammation and Alzheimer’s disease pathology

**DOI:** 10.1101/2021.05.30.446344

**Authors:** Zhengrong Zhang, Hana Na, Qini Gan, Qiushan Tao, Yuriy Alekseyev, Junming Hu, Zili Yan, Jack B. Yang, Hua Tian, Shenyu Zhu, Qiang li, Ibraheem M. Rajab, Jan Krizysztof Blusztajn, Benjamin Wolozin, Andrew Emili, Xiaoling Zhang, Thor Stein, Lawrence A. Potempa, Wei Qiao Qiu

**Author notes:** Corresponding author: Wendy Wei Qiao Qiu, M.D., Ph.D. Department of Psychiatry Department of Pharmacology & Experimental Therapeutics Boston University School of Medicine 72 East Concord Street, R-623 Boston, MA 02118.

## Abstract

**BACKGROUND:** C-reactive protein (CRP) in peripheral inflammation is associated with increased Alzheimer’s disease (AD) risk in Apolipoprotein E4 (ApoE4), but not ApoE3 or E2, humans. It remains unknown whether peripheral monomeric CRP (mCRP) induces AD pathogenesis through some receptor of blood-facing endothelia in the brain in an ApoE genotype dependent fashion.

**METHODS:** We used human samples, ApoE knock-in and deficient mouse models, and primary brain endothelia. Different ApoE mice were intraperitoneally (i.p.) injected with mCRP. The characterizations by immunostaining, proximity ligation assay (PLA) and siRNA were conducted to identify the receptor for mCRP. Brain microvessel and endothelia were isolated for RNA sequencing to explore the molecular pathway.

**RESULTS:** We demonstrate that CD31 (PECAM-1), a blood-facing endothelial receptor in brain, is a competitive target of both mCRP and ApoE protein. ApoE2 competes more strongly with mCRP for CD31 than ApoE4 does, and expressing ApoE4 or knocking out ApoE gene results in higher levels of mCRP-CD31 binding, leading to a decrease of CD31 expression but an increase in CD31 phosphorylation, along with greater cerebrovascular damage and AD pathology. This competitive binding mediates differential endothelial molecular responses depending on ApoE genotype, increasing cerebrovascular inflammation and mitochondria impairment in ApoE4 mice, while inducing vasculogenesis and protective changes in the presence of ApoE2.

**CONCLUSIONS:** Our study reveals a novel and dynamic endothelial ApoE-mCRP-CD31 pathway for AD pathogenesis during chronic inflammation and provides some insight into the opposing ApoE4-neurodegenerative and ApoE2-neuroprotective effects in AD.

**Clinical Perspective:** *WHAT IS NEW?:*

- CD31 is a competitive target of both mCRP and ApoE in brain endothelia in an ApoE-allele dependent pattern
- mCRP increases CD31 phosphorylation in the brain endothelia and damages cerebrovasculature in ApoE4 carriers and AD brains
- mCRP expression results in neuroprotective or neurodegenerative pathway activation in an ApoE-dependent manner

*WHAT ARE THE CLINICAL IMPLICATIONS?:*

- Although ApoE4 is a major genetic risk factor of AD, some ApoE4 carriers do not develop AD by the age of 90.
- Elderly people often experience peripheral inflammatory attacks and develop chronic low-grade inflammation, which results in the formation and release of mCRP. Because CRP is routine clinical laboratory test, clinicians can use blood CRP level to predict AD risk in ApoE4 carriers.
- Evidence of Apoe4 genotype and chronic low-grade inflammation stages marked by elevated CRP levels should be targeted in personalized treatment and clinical trials for AD.

---

Peripheral chronic inflammation is associated with increase Alzheimer’s disease (AD) risk ^1^. C-reactive protein (CRP) plays a major role in peripheral inflammation with two forms in the body: 1) native pentameric CRP (pCRP) oligoprotein is produced during active inflammatory reactions ^2^; and 2) monomeric CRP (mCRP) is produced during the chronic phase by the irreversible dissociation of pCRP; mCRP has a much lower aqueous solubility than pCRP and is known to cause tissue damage ^3^. Our recent human study found that elevated CRP in blood is associated with an increased risk of AD in Apolipoprotein E4 (ApoE4), but not in ApoE3 or ApoE2, human carriers ^4^. Preclinical study shows that direct injection of mCRP into the hippocampus of 3xTg AD mice enhances the severity of AD-like pathology in the brain of 3xTg AD model mice ^5^.

The blood-brain-barrier (BBB) of the cerebrovasculature is composed of both brain-facing and blood-facing cells. Because of the existence of the BBB, most blood-based inflammatory factors cannot or barely penetrate into the brain. However, almost all of these factors, including CRP, are exposed to blood-facing endothelia in the BBB. Thus, it is possible for blood CRP targeting blood-facing endothelia and causing brain AD pathology. In addition, ApoE isoforms in blood are also exposed to brain endothelia. It is unclear whether ApoE4 affects blood-facing cell, e.g endothelia, of the BBB. The cerebrovasculature is known to be altered in the AD brain ^6^. We therefore set out to used ApoE knock-in plus ApoE^-/-^ mice and both *in vivo* and *in vitro* analyses as well as human brain tissues to understand whether and how CRP and ApoE4 may interact to affect AD pathogenesis in the brain in an ApoE allele-specific fashion.

In this study, after elevating peripheral mCRP in the mice, we find that mCRP mainly bound to brain endothelia, but not to brain neurons or glia, in the brain of ApoE4 knock-in mice. This prompts us to search for some endothelial cell receptor(s) that interact with blood mCRP and may also be regulated by ApoE isoforms to cause cerebrovascular neuroinflammation in AD. We identify PECAM-1 (CD31) as an endothelial receptor for peripheral mCRP, that induces phosphorylation of CD31 and subsequent neurovascular damage. Importantly, we find that mCRP-CD31 binding is antagonized by ApoE in an allele-specific fashion, with ApoE2 competing most strongly for CD31, ApoE3 somewhat less and ApoE4 not competing well. CD31 is a cellular adhesion and signaling receptor comprising six extracellular immunoglobulin-like homology domains, a transmembrane domain and a cytoplasmic domain that becomes serine and tyrosine phosphorylated upon cellular activation to regulate vascular inflammation ^7, 8^. This differential binding of ApoE to CD31 mediates differential endothelial responses, causing an increase in cerebrovascular inflammation and AD pathogenesis in ApoE4 animals while inducing vasculogenesis and protective changes in the presence of ApoE2.

## Methods

For all the quantification measurements described below, the researchers who directly conducted counting and other measurements were blinded to ApoE genotypes, treatments and AD diagnosis.

### Human FHS ApoE and CRP data

Human serum CRP concentrations were obtained from the existing data of the Framingham Heart Study (FHS) offspring cohort (Gen 2). The source population comprised 3239 participants who met the following criteria: 1) were aged 30 years or older (range: 33 to 89 years, mean: 61.1 ± 9.5 years) at the time of the 7^th^ health exam (1998-2001), and 2) consented to use their genetic information (i.e., ApoE genotype). Human ApoE genotype were divided into three groups: ApoE2 (ApoE2/2 or 2/3, n = 463), ApoE3 (ApoE3/3, n = 2076), and ApoE4 (ApoE3/4 or 4/4, n = 665). Those with ApoE2/4 (n = 62) were excluded in the analysis.

### Mice and experimental treatments

Human ApoE genetic knock-in mice and ApoE knockout mice were purchased from Taconic Biosciences, Inc. (APOE2: #1547-F, APOE3: #1548-F, APOE4: #1549-F, ApoE^-/-^: Rensselaer, NY, USA). The human ApoE gene (either the ApoE2, ApoE3 or ApoE4 allele) replaces the endogenous mouse ApoE gene in these mice. C57BL/6 wild-type mice were purchased from Jackson Laboratory (#000664, Bar Harbor, ME, USA) to be used as a control group.

All mice were maintained in microisolator housing in the animal facility at Boston University School of Medicine. Recombinant mCRP was produced as described ^9^. Female ApoE mice aged 9-11 months were intraperitoneally (i.p.) injected with mCRP three days per week (Monday, Wednesday and Friday) for 6 weeks. mCRP was dissolved in phosphate buffered saline (PBS) before injection. Vehicle-treated mice were injected with PBS only as a control (n = 15-18 mice in each examined condition). All animal procedures were performed in accordance with the National Institutes of Health Guide for the Care and Use of Laboratory Animals and were approved by the Boston University Animal Care and Use Committee.

### Isolation, culture, and characterization of CD31^+^ brain endothelial cells (BECs)

Brain tissue samples including the cortex and hippocampus obtained from experimental mice that did or did not receive the mCRP treatment were gently dissociated into single-cell suspensions using the Adult Brain Dissociation kit (#130107677, Miltenyi Biotec, Auburn, CA, USA). Single-cell isolation and characterization of CD31^+^ cells were performed as previously described with a minor modification ^10^. Mice were deeply anesthetized by isoflurane and perfused by cold PBS. Brains were removed and dissected into the cortex and hippocampus using forceps in sterile conditions. The tissues were cut into eight slices with a razor blade and dissociated into a cell suspension using enzymatic buffer. Briefly, the tissues were enzymatically digested with the components for the mechanical dissociation step in the gentleMACS™ Octo Heat Dissociator. Following dissociation, myelin and cell debris were removed using the Debris Removal Solution. The procedure was followed by subsequent removal of erythrocytes using the Red Blood Cell Removal Solution. At this stage, we applied two methods: 1) flow cytometry to characterize the molecular signature of the cells; and 2) CD31 microbead-based isolation and culture of BECs.

1. Flow cytometry: The cell pellet was resuspended in FACS buffer (0.5% BSA, 2 mM EDTA in PBS) and labeled with anti-mouse CD31-PE (1:50, #130-111-354, Miltenyi Biotec, Auburn, CA, USA), anti-mouse CD45-FITC (1:50, #130-116-500, Miltenyi Biotec, Auburn, CA, USA) and NIR for exclusion of dead cells (1:10^3^, #425301, BioLegend, San Diego, CA, USA). Using flow cytometry (BD Bioscience), the cell samples in 0.5 ml FACS buffer were separated into different cell populations, and CD31^+^ BECs were sorted in bulk for further experiments.
2. Microbeads Endothelial cells were enriched by depletion of CD45^+^ cells with CD45 microbeads (#130-052-301, Miltenyi Biotec, Auburn, CA, USA) followed by positive selection using CD31 microbeads (#130-097-418, Miltenyi Biotec, Auburn, CA, USA) in the magnetic separator. CD31^+^ BECs were resuspended in fresh EBM-2 basal medium with all supplements (#CC-3202, EGM™-2-MV BulletKit™, Lonza, Portsmouth, NH, USA). Then, the BECs were seeded in 96-well plates coated with collagen type I (5 µg/mL, #354231, BD Bioscience, San Jose, CA, USA) at a density of 10^4^ cells per well. The medium was changed every 2 days.

On day 5, WT brain endothelial cells were treated *in vitro* with different concentrations of mCRP or vehicle control and incubated for different periods of time up to 24 hours (hrs). To explore the effects of different ApoE protein isoforms on mCRP, recombinant ApoE2, ApoE3 or ApoE4 (Perotech, Inc., Cranbury, NJ, USA) was added to CD31^+^ BECs at final concentrations of 0.03 to 3 µM and incubated for 1 hr before mCRP was added at a final concentration of 10 µg/mL. The experimental cells were fixed and processed for ApoE or mCRP and CD31 colocalization analysis using the proximity ligation assay (PLA) approach. Cells were incubated with primary antibodies (anti-phos-CD31, anti-CD31, and anti-mCRP) and subsequently stained with secondary antibodies (Invitrogen, Carlsbad, CA, USA).

### Immunofluorescence characterization

Immunofluorescence was used to characterize the postmortem human- and mouse brains. Mouse brains were collected after PBS perfusion, post-fixed in 4% paraformaldehyde for 48 hours, and changed to 30% sucrose in PBS at 4°C. Coronal cryosections (30 ìm in thickness) were used for the free-floating staining method. For frozen human postmortem brain, the sample was embedded in OCT compound, cut into 16 µm thick cryosections and mounted on gelatin-coated histological slides. The sections were allowed to air dry for 30 min and immediately fixed in ice-cold fixation buffer for 15 min. Brain slides were preincubated in blocking solution with 5% [vol/vol] horse serum (Sigma-Aldrich, St Louis, MO, USA) in 1× Tris-buffered saline for 2 hours at room temperature. The slides were incubated individually with primary antibodies overnight. Brain slides were then stained with secondary antibodies conjugated with Alexa Fluor 488, 594, 549, or 647 (1:500, Thermo Fisher Scientific, Carlsbad, CA, USA) for 1 hour at room temperature. The sections were mounted with ProLong Gold antifade reagent with DAPI for nuclear staining (#P36935, Thermo Fisher Scientific, Carlsbad, CA, USA). The stained slides were observed under fluorescence microscopy (Carl Zeiss, Germany).

The following primary antibodies were used: 1) 3H12 antibody (1:50) against mCRP ^11^; 2) anti-CD31 antibodies (#550274, BD Biosciences, San Jose, CA, USA; #ab28364, Abcam, Cambridge, MA, USA; #BBA7, R&D systems, Minneapolis, MN, USA); 3) anti-phosphorylated CD31 at tyrosine 702 (pCD31) antibody (#ab62169, Abcam, Cambridge, MA, USA); 4) anti-ApoE antibody (#701241, Invitrogen, Carlsbad, CA, USA); 5) anti-phosphorylated tau (pTau) PHF1 antibody (1:200); 6) anti-NeuN antibody (#ab177487, Abcam, Cambridge, MA, USA) to identify neurons; 7) anti-GFAP antibody (#14-9892-82, Fisher Scientific, Hampton, NH, USA) for an astrocyte biomarker; 8) anti-CD68 (#MCA1957GA, Bio-Rad Laboratories, Hercules, CA, USA) and anti-Iba-1 (#019-19741, Wako Chemicals, Osaka, Japan) antibodies for microglia biomarkers; 9) anti-Von Willebrand Factor antibody (#ab11713, Abcam, Cambridge, MA, USA) to identify vascular damage; 10) anti-CD3 (#ab16669, Abcam, Cambridge, MA, USA; #MAB4841, R&D systems, Minneapolis, MN, USA) and anti-CD8 (#NBP1-49045SS, Novus Biologicals, Littleton, CO, USA) antibodies to identify T lymphocytes; 11) anti-CD19 antibody (#NBP2-25196SS, Novus Biologicals, Littleton, CO, USA) to identify B lymphocytes; and 12) anti-CD14 antibody (#11-0141-82, Invitrogen, Carlsbad, CA, USA) to detect monocytes; 13) anti-Lectin antibody (#DL-1174, Vector, Burlingame, CA, USA) and anti-CD144 antibody (#14-1441-82 Invitrogen, Carlsbad, CA, USA) to label vascular and endothelia.

To evaluate immunostaining results, ImageJ was used to measure total intensity after adjusting the threshold. The data obtained from two independent researchers who were blinded to the treatment groups were pooled and averaged.

### Proximity Ligation Assay

A proximity ligation assay (PLA) was applied to both frozen brain sections and primary endothelial cells after fixation to investigate protein-protein interactions ^12^. The samples were washed with PBS and incubated with blocking buffer at 37°C for 1 hour. To detect mCRP and CD31 interaction, samples were incubated overnight at 4°C with mouse anti-mCRP antibody (1:50, 3H12) and either goat anti-CD31 antibody (1:200, #AF3628, R&D systems, Minneapolis, MN, USA) for mouse samples or rabbit anti-CD31 antibody (1:200, #ab28364, Abcam, Cambridge, MA, USA) for human samples. To detect ApoE and CD31 interaction, samples were incubated overnight at 4°C with rabbit anti-ApoE antibody (1:200, #701241, Invitrogen, Carlsbad, CA, USA) and either rat anti-CD31 antibody (1:200, #550274, BD Biosciences, San Jose, CA, USA) for mouse samples or mouse anti-CD31 antibody (1:200, #BBA7, R&D systems, Minneapolis, MN, USA) for human samples. The samples were similarly incubated with mouse IgG and goat IgG or rabbit IgG and rat IgG as control groups, respectively. Proximity ligation was then conducted in situ as described by the manufacturer’s instructions (#DUO92007, Sigma-Aldrich, St Louis, MO, USA). We used the Duolink PLA probes anti-mouse PLUS and anti-goat MINUS to visualize mCRP/CD31 interactions and the Duolink PLA probes anti-mouse PLUS and anti-rabbit MINUS to visualize ApoE/CD31 interactions by Duolink in Situ Detection Reagents Orange. The samples were further incubated with Alexa Fluor 488- and 647-conjugated secondary antibodies (Invitrogen, Carlsbad, CA, USA) for 1 hour at room temperature. Following serial washes, the samples were stained with DAPI and observed with a fluorescence microscope (ZEISS Axio Observer).

### Microvessel isolation

Microvascular of mouse brain were isolated using dextran gradient centrifugation followed by sequential cell strainer filtrations as previously described^13^. Briefly, brains were carefully remove and transfer immediately to cold MCDB131 medium. The cortex and hippocampus were dissected and all visible white matter was discarded. The isolated tissue was placed to MCDB131 medium and homogenized using a loose fit 7ml Dounce tissue grinder. The samples were transferred into 14ml round-bottom centrifuge tube and then centrifuged at 2,000 g for 5 min. The pellets were resuspended in 15 % dextran/PBS (70kDa, Sigma). The samples were thoroughly mixed and centrifuged at 10,000 g for 15 mins, 4°C. The microvessel -containing pellet located at the bottom of the tubes was collected, and sequentially resuspended using 1 ml PBS. The pellet was transfer onto a 40µm cell strainer (BD Falcon, San Jose, CA, USA) and wash through <10 ml PBS. The microvascular remaining on top were collected in MCDB 131 medium containing 0.5% endotoxin-, fatty acid- and protease free BSA and fixed by 4% PFA for further fluorescent staining analysis.

### CD31 self-interfering RNA (siRNA)

Primary mouse BECs were incubated with 1µM Accell CD31-targeing siRNA (SMART pool # E-048240) and Non-targeting controls (# D-001910) for 72 h in Accell delivery media (# B-005000). Removing medium, the BECs were treated with mCRP (10 µg/ mL)/ EBM-2 basal medium for 10 h and fixed for further staining analysis.

### Microvascular profile measurements

The lengths of CD31-positive cerebrovascular profiles were measured as previously reported ^14^. Fixed brain sections were prepared and stained as described in the “Immunofluorescence characterization” section, followed by incubation with anti-CD31 primary antibody (1:200, #550274, BD Biosciences, San Jose, CA, USA; #BBA7, R&D systems, Minneapolis, MN, USA) and Alexa Fluor 594-conjugated secondary antibody.

In mouse brain sections, the CD31^+^ microvessels (vessels < 6 µm in diameter) were selected, and their lengths were measured from the 3D structures by using the ImageJ in length analysis tool in 5 randomly selected areas in the cortex. Z-stacks were collected at 1-µm steps for a total imaging depth of 16 µm. The average length of CD31^+^ microvessels was expressed as µm per mm^3^ of brain tissue.

In human brain sections, we measured the length of CD31^+^ from 3D images as above and also counted the number of vessel terminals in 2D images. The lengths of blood vessels in 2D images could be affected by either vessel breakage or distortion. CD31^+^ microvessels (length >20 µm, diameter <6 µm) were selected and counted from 5 randomly selected areas for each individual sample. The average numbers were recorded per mm^2^ of brain tissue.

### Western blots

The mouse cortex/hippocampus and postmortem frontal cortex were lysed in cold RIPA buffer supplemented with a protease and phosphatase inhibitor mixture (#78445 Thermo Fisher Scientific, Carlsbad, CA, USA). For each western blot, 40 µg of protein extract per sample was used. All samples were electrophoresed on NUPAGE 4-12% Tris-Glycine Gels (#XP04205BOX, Invitrogen, Carlsbad, CA, USA) and transferred to PVDF membranes. The membranes were blocked with 5% milk and incubated with primary antibodies. The following primary antibodies were utilized for immunoblots: goat anti-CD31 antibody (1:500, #AF3628, R&D Systems, Minneapolis, MN, USA) for mouse, rabbit anti-CD31 antibody (1:500, #ab28364, Abcam, Cambridge, MA, USA) for human, rabbit anti p-CD31 (phospho Tyr 702) antibody (1:500, #ab62169, Abcam, Cambridge, MA, USA), rabbit anti p-eNOS (Ser1177) antibody (1:500, #9571s, Cell Signaling Technology, Beverly, MA, USA), rabbit anti-eNOS antibody (1:500, #32027, Cell Signaling Technology, Beverly, MA, USA), rabbit anti NF-κB p65 antibody (1:500, #8242, Cell Signaling Technology, Beverly, MA, USA), and rat anti-CD144 (VE-cadherin) antibody (1:500, #14144182, Invitrogen, Carlsbad, CA, USA). After washing with Tris-buffered saline (TBS) with Tween 20, the membranes were incubated with the appropriate HRP-conjugated secondary antibodies in blocking buffer and imaged using a Bio-Rad imaging station. For the quantification analysis of western blots, a rectangular area was selected enclosing the bands of interest, and the intensity of each band was measured from the area showing the most intense signal using ImageJ. The signal intensities of all protein bands were normalized by the β-actin band, which served as the control in this experiment.

### Enzyme-linked immunosorbent assay (ELISA)

Proteins were extracted from the cortex, hippocampus, or cells using RIPA buffer with a protease inhibitor mixture. Venous blood was collected by cardiac puncture and placed at room temperature for 1 h. The samples were centrifuged for 15 min at 3000 rpm to obtain serum. ELISA assays were conducted for different kinds of samples. The Aβ42 (ELISA kit, **#** KMB3441, Invitrogen, Carlsbad, CA, USA) and CRP (Quantikine ELISA kit, #MCRP00, R&D systems, Minneapolis, MN, USA) levels in mouse brain extracts and serum were measured by following the manufacturer’s instructions. All assays were performed in duplicate, and the average of two values was used for further analysis.

### RNA isolation and sequencing

Total RNA was isolated from dissected CD31^+^ BECs from the mouse brain using an RNA extraction kit (#74104, Qiagen, MD, USA). The extracted RNA was stored in RNAse-free water at -80°C until transfer to the Boston University School of Medicine Microarray Core Facility. The RNA quality of the samples was assessed using the Agilent 2100 Bioanalyzer (Agilent Technologies). All samples passed a quality control threshold to proceed to library preparations and RNA sequencing. The libraries were prepared from total RNA enriched for mRNA using the NEBNext Poly(A) mRNA Magnetic Isolation Module and the NEBNext Ultra II Directional RNA Library Preparation Kit for Illumina (New England Biolabs, MA, USA) and sequenced on an Illumina NextSeq 500 instrument (Illumina, San Diego, CA, USA). Raw reads from each library were mapped against the mouse genome build mm10 using STAR (version 2.6.0c) as described in the literature ^15^. FASTQ quality was assessed using FastQC (version 0.11.7), and alignment quality was assessed using RSeQC (version 3.0.0).

### Differential gene expression and pathway enrichment analyses

Variance-stabilizing transformation (VST) was accomplished using the variance stabilizing transformation function in the DESeq2 R package (version 1.23.10). Differential expression was assessed using ANOVA and Wald tests as implemented in the DESeq2 R package. Correction for multiple hypothesis testing was accomplished using the Benjamini-Hochberg false discovery rate (FDR). Human homologs of mouse genes were identified using HomoloGene (version 68). All analyses were performed using the R environment for statistical computing (version 3.6.0).

Gene Set Enrichment Analysis (GSEA) (version 2.2.1) ^16^ was used to identify biological terms, pathways and processes that were coordinately up- or downregulated within each pairwise comparison. The Entrez Gene identifiers of the human homologs of all genes in the Ensembl Gene annotation were ranked by the Wald statistic computed between mCRP and PBS within each ApoE genotype group. Ensembl human genes matching multiple mouse Entrez Gene identifiers and mouse genes with multiple human homologs (or vice versa) were excluded prior to ranking so that the ranked list represents only those human Entrez Gene IDs that match exactly one mouse Ensembl Gene. Each ranked gene list was then used to perform preranked GSEA analyses (default parameters with random seed 1234) using the Entrez Gene versions of the Hallmark, BioCarta, KEGG, Reactome, PID, transcription factor motif, microRNA motif, and Gene Ontology (GO) gene sets in the Molecular Signatures Database (MSigDB) version 7.1 ^17^. All of the gene sets were used for analysis to facilitate full reproducibility of results. We also obtained the human single cell RNA dataset from published study ^18^ and used GSEA analyses to study gene expressions in endothelia cells in AD as described above.

### Human brain tissue and characterization

Brain temporal lobe tissues from 8 cognitive controls and 10 AD patients were obtained from the Brain Bank of the Boston University Alzheimer’s Disease Center (BU ADC). The clinical features and ApoE genotypes are listed in Supplement Table 1. Immunostaining and western blots as described above were used to characterize the levels of mCRP, the binding of mCRP-CD31 or ApoE-CD31, and the levels of pCD31 in the brain tissues. The lengths of CD31^+^ microvessl were measured from 3D imaging and the terminal ends of CD31^+^ microvessel were counted from 2D imaging. This process was conducted by researchers blinded to the diagnoses.

### Quantification and statistical analysis

Experimenters were blinded to the genotypes during testing. All data are presented as the mean ± standard error of the mean (SEM). Statistical analyses were performed using GraphPad Prism (version 8.0). For mouse data, datasets were analyzed for significance using Student’s unpaired two-tailed t test for two groups. Comparisons between three or more groups were conducted using one-way or two-way analysis of variance (ANOVA). Post hoc multiple comparisons were carried out using Tukey’s test. For human data, we performed univariate analyses stratified by ApoE genotype (ApoE2-2/2 or 2/3; ApoE3-3/3; ApoE4-3/4 or 4/4) or control vs. AD brains. Mean ± SD was determined and ANOVA tests were conducted on variables. The correlation analysis was conducted by Pearson or Spearman test according to the distribution pattern of the data. Statistical significance was defined as p value < 0.05.

## Results

### mCPR binds to CD31 of brain endothelia in an ApoE-allele dependent pattern

To address if peripheral mCRP and ApoE in blood act on blood-facing endothelia in the brain, we examined human brain tissue samples and mice with different ApoE genotypes. Both human and mice consistently showed different concentrations of serum CRP following the pattern ApoE2>ApoE3>ApoE4 (Supplement Figure 1A). Furthermore, we conducted i.p. injections of human pCRP or mCRP into ApoE knock-in mice and found more CRP, particularly mCRP, deposited in mouse ApoE4 brain than in the ApoE3 and ApoE2 brains (Supplement Figure 1B). Chronic elevation of peripheral mCRP levels by i.p. injection (200 µg/kg) three times per week for 6 weeks (n = 15-18 in each group) produced the highest levels of mCRP in the blood of ApoE2 mice; in contrast, the highest levels of mCRP in the brain were observed in ApoE4 mice (Supplement Figure 1C and 1D). In addition, brain mCRP and blood mCRP were negatively associated (Supplement Figure 1C). These data suggest that peripheral mCRP proteins gain access to the brain in an ApoE-allele specific fashion, with the most appearing in the ApoE4 brains.

**Figure 1.**
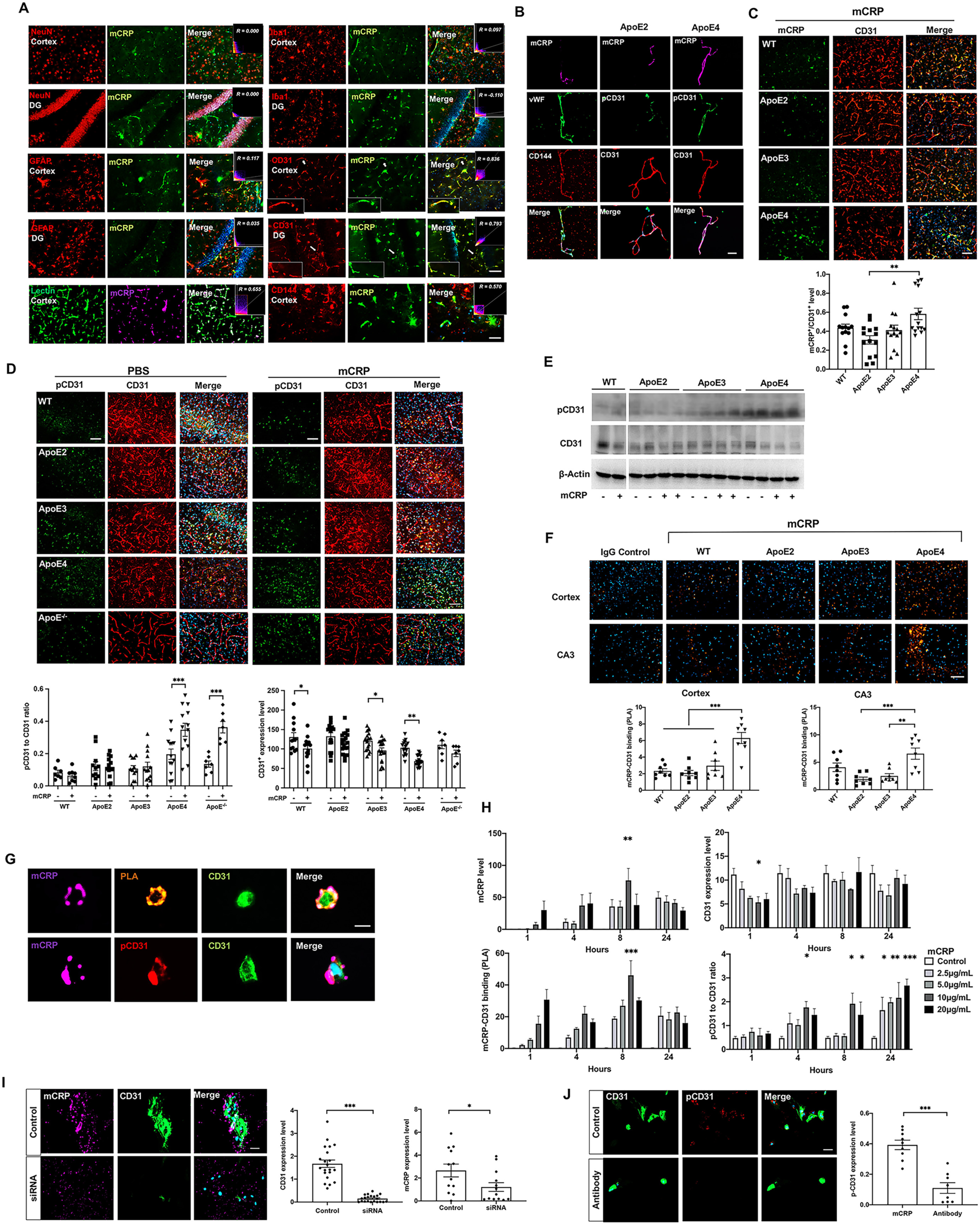
Characterization of peripheral mCRP in blood-facing endothelial based on ApoE genotype-specific regulation. 1A. To determine which cell type binds mCRP in the brain after the intraperitoneal (i.p.). injection, double immunostaining of mCRP and different types of cell markers was conducted in the cortex and hippocampal DG region. Cell markers, including the neuronal marker NeuN, the microglial marker Iba1, the astrocyte marker GFAP, vascular marker Lectin and the endothelial cell marker CD31 and CD144, were applied individually. Correlation coefficients and R values were calculated for each protein pair detected and are shown. 1B Immunofluorescence analysis of isolated brain microvessels from mice treated with mCRP (purple) were stained with antibodies for endothelial cell components CD144 (red), vWF (green), CD31 (red) and its phosphorylation pCD31 (green). Nuclear stained with DAPI. The bar is 20 µm. 1C. Representative images of cortex stained with CD31 (red) and mCRP (green) and merged (yellow) are shown to observe mCRP deposits in CD31 positive regions. The yellow fluorescence intensities were quantified and compared among different genotype mice with i.p. mCRP treatment. *p* = 0.0016, n = 11-14 mice in each group. 1D. Representative images of double immunostaining of phos-CD31 (pCD31, green) and CD31 (red) and the merged images (yellow), nuclei stained with DAPI, on the cortex of WT mice, ApoE^-/-^ mice and mice expressing different ApoE genotypes after i.p. treatment with PBS vs. mCRP are shown. Total CD31 and pCD31 were quantified by fluorescence intensity and the pCD31/CD31 ratio (ApoE4 *p* < 0.001, ApoE^-/-^ *p* < 0.001) total CD31 (WT *p* = 0.03, ApoE3 *p* = 0.03, ApoE4 *p* = 0.001), n = 7-19 in each group. 1E. Western blots showed that mCRP increased the level of pCD31 and decreased the CD31 expression levels in the hippocampal region in ApoE4 mice but not in ApoE3 or ApoE2 mice. 1F. Proximity ligation assay (PLA) was performed on the cortex (upper panel) and hippocampal CA3 region (lower panel) to examine the binding of mCRP and CD31 in WT mice and mice expressing different ApoE genotypes after i.p. injection of PBS vs. mCRP. Positive PLA fluorescence signals are shown in orange; nuclei were stained with DAPI (blue). Quantifications of orange fluorescence were conducted and are shown: WT vs. ApoE4 *p* < 0.0001, ApoE2 vs. ApoE4 *p* < 0.0001, and ApoE3 vs. ApoE4 *p* = 0.0002 for cortex; ApoE2 vs. ApoE4 *p* = 0.0004 and ApoE3 vs. ApoE4 *p* = 0.0018 for CA3. n = 8 in each group. 1G. Representative images of primary CD31^+^ BECs from WT mice treated with mCRP *in vitro* at day 5 are shown. PLA (orange) was performed to detect colocalization/binding between mCRP (purple) and CD31 (green). pCD31 (red) was detected by a specific antibody, and the nuclei were stained with DAPI. The scale bar is 10 µm. 1H. To determine whether the direct effects of mCRP on endothelia are dose-dependent and/or time-dependent, primary CD31^+^ BECs were treated with various concentrations of mCRP for 1, 4, 8 and 24 h. Quantification is shown of the mCRP deposits on the surface of cells (upper left panel), the binding affinity of mCRP with CD31 (lower left panel), the CD31 expression level (upper right panel) and the ratio of pCD31 normalized against total CD31 (lower right panel) under different concentrations of mCRP and time course. mCRP (10 µg/ml) showed maximum binding to CD31 after 8 hours of incubation (*p* = 0.0002). mCRP decreased the expression of CD31 within the first hour of incubation (*p* = 0.04) and increased the levels of pCD31 after incubation for up to 24 hours (*p* < 0.001) in a dose-dependent manner. At least three independent experiments were conducted for each condition. 1I. Primary BECs were transfected with CD31-targeting siRNA to effectively knockdown CD31 expression (*p* < 0.001) before adding mCRP (10 µg/mL) stimulation. Fluorescence immunostainings with antibodies, mCRP (purple), CD31 (green) and pCD31 (red) were shown. Nuclear was stained with DAPI. The quantifications were shown that compared to control cells, BECs after silencing CD31 with siRNA had lower mCRP binding (*p* = 0.04). 1J. Primary BECs were incubated with CD31-specific antibody followed by adding mCRP (10 µg/mL) stimulation. A significant decreased pCD31 levels were observed in BECs after blocking CD31 by the antibodies (*p* < 0.001). Data are shown as the mean ± SEM. One or two -way ANOVA with Tukey’s post hoc test and Pearson or Spearman correlation test were applied. \**p* < 0.05, \*\**p* < 0.01, \*\*\**p* < 0.001, \*\*\*\**p* < 0.0001. The scale bar is 50 µm.

To investigate which brain cells mCRP binds to, we used antibodies against mCRP and biomarkers of different brain cell types. By assessing the colocalization of fluorescence-labeled proteins, we found that after i.p. injection, mCRP colocalized with endothelial cells marked by CD31 but not with the markers for neurons, astrocytes or microglia in the mouse brain (Figure 1A). To further confirm that mCRP binds to endothelia, we used two other endothelia markers, lectin and CD144 and found that mCRP deposited in Lectin-positive blood vascular, especially co-localized with endothelial cells marked by CD144 and CD31 (Figure 1A). Further, we isolated the microvessel and used endothelial related markers, von Willebrand factor (vWF) and CD144, to confirm that mCRP did bind to brain endothelia (Figure 1B).

Using ApoE knock-in mice, we found elevated peripheral mCRP deposits in CD31 positive endothelia regions in the ApoE4 brain compared with the ApoE2 brain (Figure 1C) and their isolated microvessels (Figure 1B, right two columns). Using both immunostaining (Figure 1D) and western blot (Figure 1E and Supplement Figure 1E), we found that mCRP reduced the expression levels of total CD31 and caused increase of CD31 phosphorylation in an ApoE allele-specific fashion with the pattern of ApoE4>ApoE3>ApoE2 in the brain that was confirmed in the isolated brain microvessels (Figure 1B, right two columns). Like ApoE4 mice, ApoE^-/-^ mice also had a significant increase in pCD31 with mCRP treatment (Figure 1D).

Since phosphorylation of the CD31 cytoplasmic domain has been shown to inhibit CD31 function ^19^, we hypothesized that peripheral mCRP binds and acts on endothelial CD31 as a receptor to affect differential phosphorylation level. To prove this, proximity ligation assay (PLA) was used to indicate that mCRP directly bound to CD31 in the cortex and hippocampus (Figure 1F) with an intensity pattern reflective of ApoE allele type (ApoE4>ApoE3>ApoE2). To further investigate if mCRP binding to endothelial CD31, brain CD31^+^ endothelial cells (BECs) were isolated from mouse brain tissues ^10^ and used to examine the effects of mCRP on CD31 in the presence of different ApoE isoforms *in vitro* (Figure 1G-J). Indeed, exogenous mCRP was colocalized with CD31, which is highly expressed on the surfaces of primary brain endothelial cells (Figure 1G).

Figure 1H shows that after adding mCRP to the cultures, the highest level of mCRP-CD31 binding was found at 8 hrs of incubation (upper left panel); CD31 expression was lowed in the first hr (upper right panel), with the highest mCRP-CD31 binding at 8hrs (lower left panel) and the production of pCD31 increasing starting at 4 hrs of incubation and reaching its highest level at 24 hrs (lower right panel) in a mCRP dose-dependent manner (Supplementary Figure 1F). Further, after CD31 was knockdown by siRNA in primary endothelia, no or minimum mCRP bound to endothelia (Figure 1I). After CD31 was blocked by incubating with the CD31-specific antibody, pCD31 level was reduced after mCRP treatment (Figure 1J). In summary, these results indicate that endothelial CD31 in the brain is a receptor for mCRP from peripheral inflammation.

### CD31 is a competitive target of both mCRP and ApoE in brain endothelia

We first examined brain levels of ApoE protein across ApoE genotypes and found that ApoE2 mice had the highest level of ApoE protein regardless the treatment (Figure 2A). To understand how ApoE genotype affects mCRP-CD31 binding and pCD31 production, we used PLA and did find that peripheral i.p. injection of mCRP significantly increased ApoE-CD31 binding only in the brains of ApoE2, but not in ApoE3 and ApoE4, mice (Figure 2B), which was in direct opposition to the mCRP-CD31 binding data (Figure 1F). We found no binding/colocalization between ApoE and mCRP (Supplement Figure 2A). These data suggest that mCRP and ApoE may differentially compete for CD31 binding.

**Figure 2.**
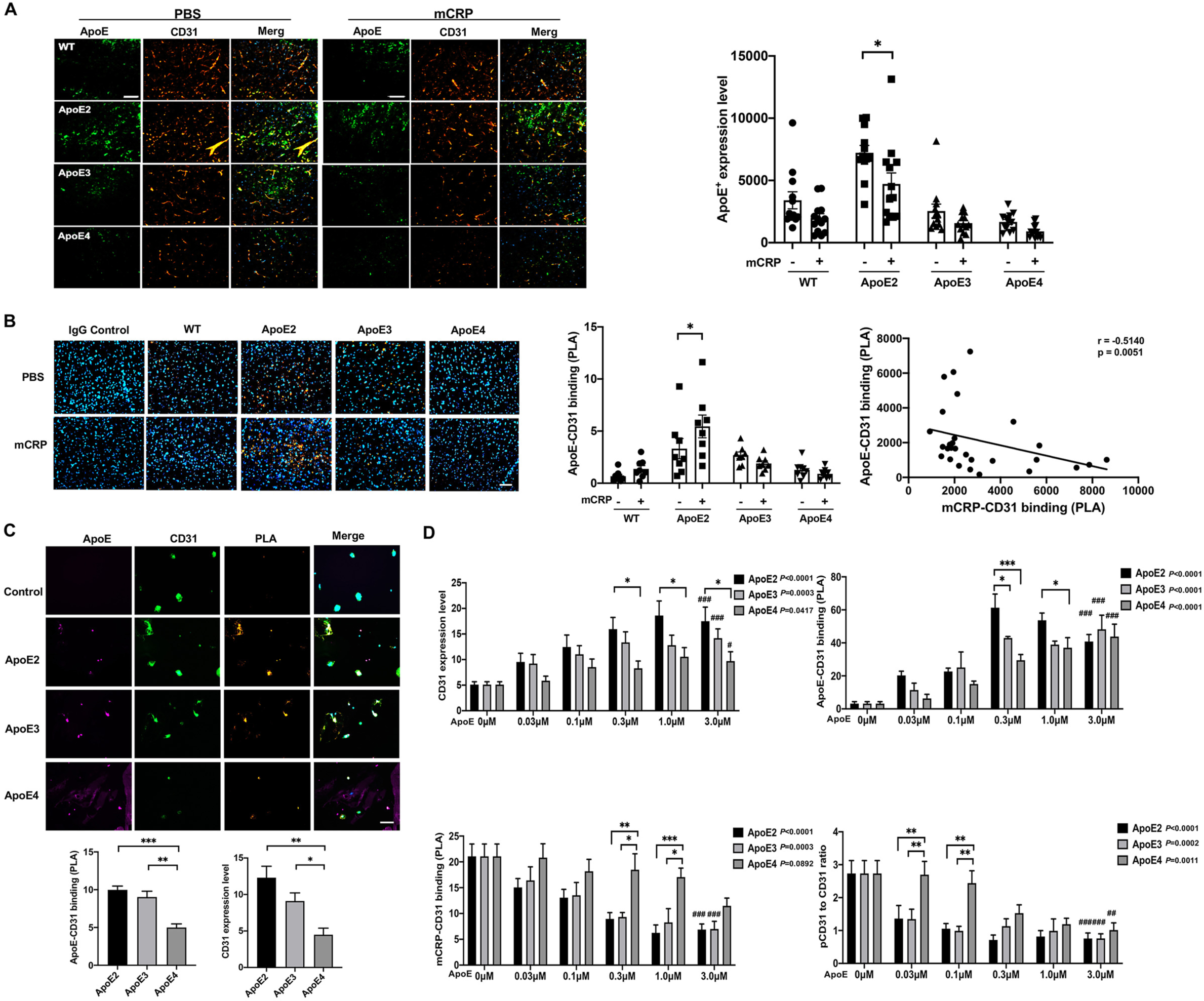
The competition binding between mCRP and ApoE to CD31^+^ endothelial cells. 2A. Representative images of double immunostaining of ApoE (green) and CD31 (red) and the merged images (yellow) of the cortex sections of WT vs. different ApoE knock-in mice are shown. The ApoE^+^ fluorescence intensity was compared between PBS and mCRP-treated mice and reached a significant difference only in the ApoE2 mice (*p* = 0.02). n = 11-14 mice in each group. 2B. PLA was applied to detect the binding of ApoE and CD31 on cortex sections from different ApoE genotype mice treated with PBS vs. mCRP. Positive PLA fluorescence signals are shown in orange, and nuclei are stained with DAPI (blue). Quantifications of orange fluorescence intensity of ApoE-CD31 binding in the cortex are shown and reached a significant difference only in ApoE2 mice (*p* = 0.04). PLA (mCRP-CD31) levels were negatively correlated with PLA (ApoE-CD31) in the brain (r = -0.51, *p* = 0.005). n = 8 in each group. 2C. Representative images of primary CD31^+^ BECs treated with 0.3 µM ApoE2, ApoE3 or ApoE4 protein are shown. The interaction/binding between ApoE (purple) and CD31 (green) was detected by PLA (orange). The positive PLA signal and CD31 protein level were quantified and compared. ApoE4 protein had the lowest binding with CD31 compared to ApoE2 (*p* = 0.0002) and ApoE3 (*p* = 0.0013) proteins. In addition, ApoE4 exhibited the most significant decrease in CD31 expression in CD31^+^ BECs compared to ApoE2 (*p* < 0.001) and ApoE3 (*p* = 0.04). The scale bar is 50 µm. 2D. To examine how different ApoE isoform proteins influence the effect of mCRP on endothelia, primary CD31^+^ BECs were preincubated with different concentrations of ApoE2, ApoE3 or ApoE4 protein for one hour followed by the addition of 10 µg/mL mCRP for 6 hours. Quantification of the CD31 expression level (upper left panel), the binding of ApoE-CD31 (upper right panel) and mCRP-CD31 (lower left panel) and the pCD31/CD31 ratio (lower right panel) were conducted. *Three ApoE isoform proteins were compared for each concentration, and the statistical significance is shown. #The comparison among different doses of protein for one isoform of ApoE; at least three independent experiments were conducted. Data are shown as the mean ± SEM. One or two -way ANOVA with Tukey’s post hoc test and Pearson correlation test were applied. \**p* < 0.05, \*\**p* < 0.01, \*\*\**p* < 0.001; #*p* < 0.05, ##*p* < 0.01, ###*p* < 0.001.

Further, we added equal concentrations of recombinant ApoE2, ApoE3 or ApoE4 to the primary BECs and incubated for one hour to confirm ApoE-CD31 binding. Figure 2C shows that endothelial ApoE-CD31 binding and CD31 expressions were the lowest in the presence of ApoE4 and the highest for ApoE2. To directly examine whether ApoE isoforms compete with and influence the mCRP effects on endothelia, pre-incubation of BECs with different ApoE isoforms and with different concentrations was followed by adding mCRP (Figure 2D). This resulted in the greatest levels of CD31 expression (upper left panel) and highest levels of ApoE-CD31 binding (upper right panel) in ApoE2-pretreated cells with the reverse observed in ApoE4-pretreated cells in a dose- and time-dependent pattern; in contrast, mCRP-CD31 binding (lower left panel) and pCD31 production (lower right panel) were increased the most in ApoE4-treated and the least in ApoE2-pretreated cells. Thus, increased ApoE-CD31 binding inhibits the mCRP-CD31 interaction and reduces the production of pCD31 in a dose-dependent manner. These findings indicated that ApoE4 protein is less likely to bind to CD31 than ApoE2 or ApoE3, and thus increases the overall levels of mCRP-CD31 and pCD31 production in endothelial cells.

### mCRP increases CD31 phosphorylation in the brain endothelia and damages cerebrovasculature in ApoE4, but not ApoE2 or ApoE3, mice

We hypothesized that mCRP may cause detrimental effects via the mCRP-CD31 interaction and the resulting decreasing in total CD31 but increase in pCD31. To assess this, we first investigated actions of peripheral mCRP on the mouse cerebrovasculature and whether these effects were sensitive to the ApoE genotype. We found that both ApoE^-/-^ and ApoE4 mice exhibited shorter cerebrovasculature lengths when stained with a CD31 antibody than ApoE2 and ApoE3 mice (Figure 3A top row). Elevating peripheral mCRP further shortened the length of the CD31^+^ microvessels with a severity pattern of ApoE^-/-^>ApoE4 > ApoE3, but not ApoE2, in the cortex (Figure 3A, bottom row) and hippocampus (Supplement Figure 3). This suggests that ApoE4 mice, equivalent to ApoE^-/-^, are more susceptible to cerebrovascular damage, with mCRP and pCD31 exacerbating these effects.

**Figure 3.**
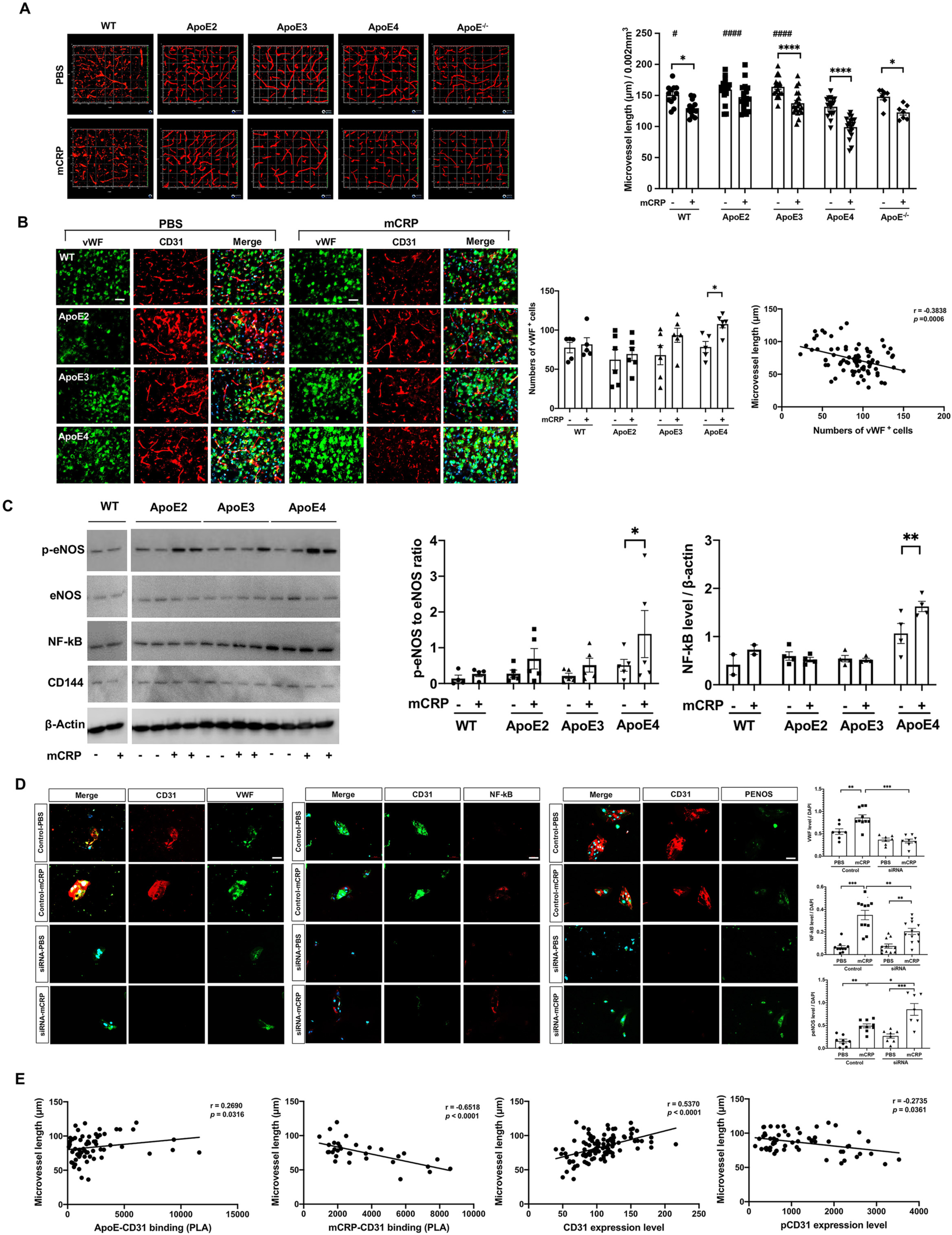
The impact of peripheral mCRP on cerebrovascular integrity in an ApoE genotype-dependent manner. 3A. After the i.p. treatment of PBS (top column) vs. mCRP (bottom columns), 3D images of cortex sections stained for CD31 (red) in WT, different ApoE-expressing and ApoE^-/-^ mice were used to examine cerebrovasculature. The length of CD31-positive microvasculature in the cortex was quantified and compared between the PBS and mCRP groups in WT (*p* = 0.01), ApoE3 (*p* < 0.0001), ApoE4 (*p* < 0.0001) and ApoE^-/-^ (*p* = 0.04) mice. # Comparison of ApoE genotypes in the PBS group, ApoE4 vs WT *p* = 0.04, ApoE2 *p* < 0.0001, ApoE3 *p* < 0.0001. n = 7-19 mice in each group. 3B. To examine the impact of elevated peripheral mCRP on cerebrovascular damage, we conducted immunostaining of a vascular damage biomarker, von Willebrand factor (vWF, green), and CD31 (red), in the cortex of WT vs. different ApoE knock-in mice. Representative images of each genotype treated with PBS vehicle (left columns) and mCRP (right columns) are shown. The numbers of vWF-positive endothelial cells in the cortex were compared among different genotype mice in the absence and presence of i.p. mCRP and showed statistical significance only in the ApoE4 group (*p* = 0.04). A negative correlation between the length of CD31^+^ microvessels and the expression of vWF in the cortex was shown (r = -0.38, *p* = 0.0006). n = 5-6 in each condition. 3C. Western blots of different inflammatory and vascular-related proteins, including phosphorylated eNOS (p-eNOS), eNOS, NF-κB and CD144, in the cortex were conducted to examine the effects of mCRP on these proteins in each ApoE genotype. Peripheral mCRP significantly increased the expression of p-eNOS (*p* = 0.04) and NF-κB (*p* = 0.006) only in ApoE4 mice. 3D. Primary BECs were transfected with CD31-targeting siRNA to effectively knockdown CD31 expression (*p* < 0.001) before adding mCRP (10 µg/mL) stimulation. Representative images of primary BECs silencing of CD31 with mCRP treatment were stained with antibodies vWF(green), NF-κB (red), p-eNOS (green). The nuclei was stained with DAPI. The quantifications were shown that lower levels of vWF (*p* < 0.001), NF-κB (*p* = 0.003) but higher level of p-eNOS (*p* = 0.005) were found in siRNA group, compared with control group after mCRP treatment. At least three-time experiments were conducted. 3E. Different correlation analyses were conducted to examine the relationship between elevated peripheral mCRP and CD31^+^ microvessels in the brain. The graphs show that the length of CD31^+^ microvessels (y-axis) was positively associated with the CD31 level (r = 0.54, *p* < 0.0001) and the levels of PLA (ApoE-CD31) (r = 0.27, *p* = 0.03) but negatively associated with the levels of PLA (mCRP-CD31) (r = -0.65, *p* < 0.0001) and pCD31 (r = - 0.27, *p* = 0.04). Data are expressed as the mean ± SEM. Two-way ANOVA with Tukey’s post hoc test and Pearson or Spearman correlation test were applied. \**p* < 0.05, \*\**p* < 0.01, \*\*\**p* < 0.001. The scale bar is 50 µm.

We employed vWF ^20^, a biomarker of cerebrovascular inflammation and damage. mCRP treatment significantly increased the level of vWF only in the brains of ApoE4 mice (Figure 3B); in addition, a correlation analysis for all the experimental mice showed that the vWF level was negatively correlated with the length of CD31^+^ microvessels (p < 0.001) (Figure 3B). mCRP treatment also increased the levels of other vascular and inflammatory factors in ApoE4 brains, such as p-eNOS and NF-κB, while mCRP tended to decrease the level of eNOS and did not influence CD144, as shown by western blots (Figure 3C). These data indicate that mCRP causes an elevated cerebrovascular inflammatory response only in ApoE4 brains, possibly indicating that ApoE2 and E3 may prevent this effect. To confirm this, CD31 in primary endothelia was knocked down by siRNA and failed to response to mCRP, resulting in little or no change of vWF and NF-κB level but increased p-eNOS level (Figure 3D). mCRP had two effects on endothelial CD31: 1) decrease in total CD31 expression and 2) increase in pCD31 (Figure 1D). Because that siRNA almost knocked out the CD31 expression (Figure 1I) so there would be no pCD31 production, it indicates that mCRP, through decreasing CD31, increased production of p-eNOS as a compensating factor.

We conducted correlation analyses across all the ApoE genotype mice used in the experiments in the absence and presence of mCRP treatment (Figure 3E). Microvessel length in the brain was positively associated with ApoE-CD31 binding (r = 0.27, p = 0.03) but negatively associated with mCRP-CD31 binding (r = -0.65, p < 0.0001). Consistently, microvessel length in the brain was positively associated with CD31 expression levels (r = 0.54, p < 0.0001) but negatively with pCD31 levels (r = -0.27, p = 0.04). The data again suggest that mCRP and ApoE competition for CD31 exerts opposing effects, with the increased mCRP-CD31 binding in ApoE4 carriers leading to an increase in mCRP-CD31 binding and pCD31 expression that results in greater cerebrovascular inflammation and damage.

### mCRP and mCRP-CD31 binding lead to AD pathogenesis in ApoE4, but not ApoE2 or ApoE3, mice

We next examined whether peripheral mCRP had a direct *in vivo* effect on the development of brain AD pathology in mice that differed among ApoE genotypes. Compared to the PBS treatment, the mCRP treatment significantly induced neuronal tauopathy, an AD hallmark, which was marked by the immunostaining of pTau with the PFH-1 antibody, in the neurons of the cortex (p < 0.01) (Figure 4A) and hippocampus (p < 0.05) (Figure 4B) in ApoE4, but not in ApoE3 or ApoE2, mice. mCRP i.p injection slightly increased the level of Aβ1-42, another pathological marker of AD, only in the ApoE4 brain (Supplement Figure 4A). Elevated peripheral mCRP also significantly increased the expression of the inflammatory microglial markers Iba1 (p < 0.001) and CD68 (p < 0.05) but had a minimal effect on the expression of the glial protein GFAP only in the cortex of ApoE4 mice (Figure 4C).

**Figure 4.**
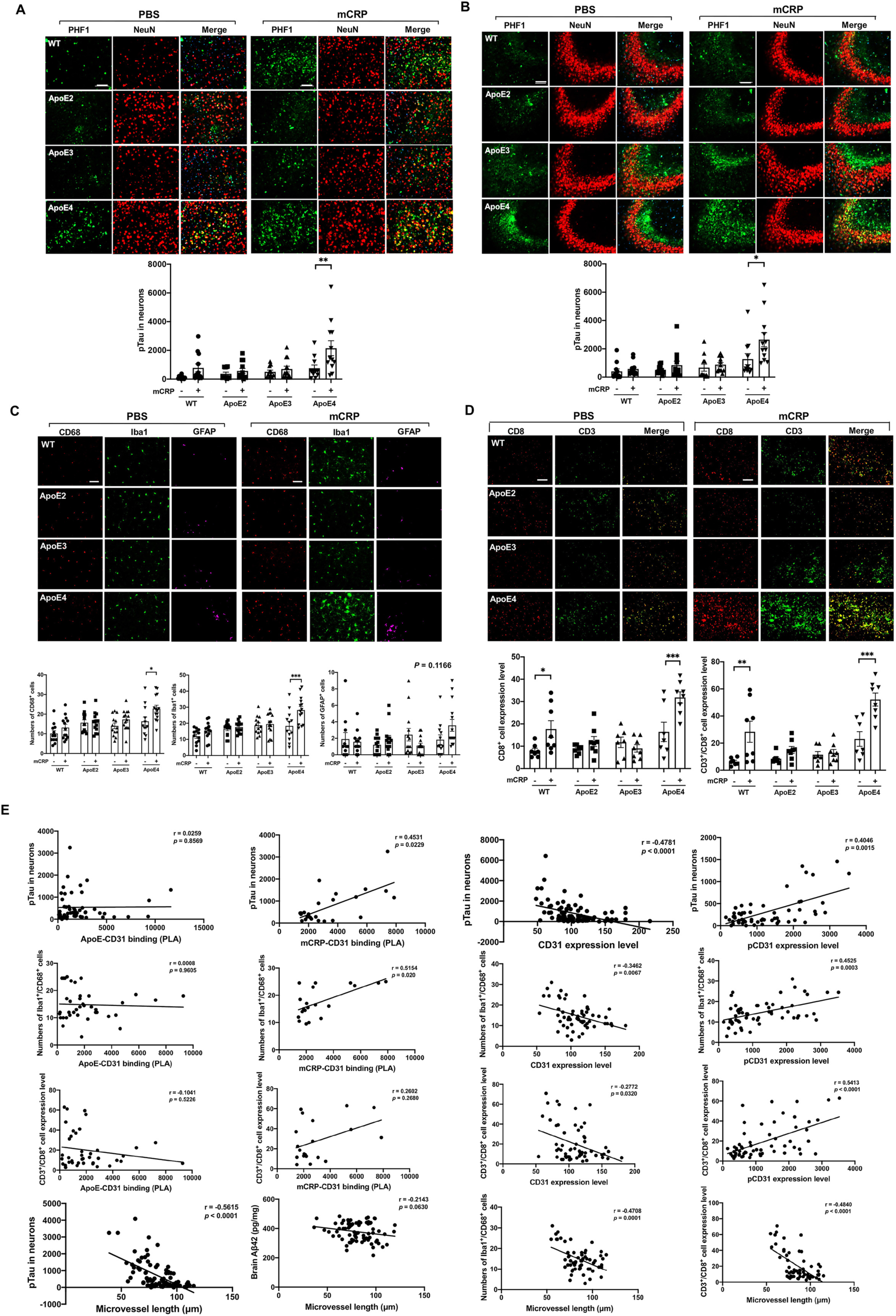
Peripheral mCRP induced neuroinflammation, extravasation of T lymphocytes and AD pathology in the ApoE4 brain. 4A and 4B. Representative images of i.p. mCRP-induced neuronal tau phosphorylation (pTau) in the cortex (A) and the CA3 region of the hippocampus (B) are shown. Double immunostaining of pTau (stained with PHF1, green) and neuronal marker (stained with NeuN, red); nuclei were stained with DAPI. Quantification of PHF1 levels in NeuN-positive cells showed statistical significance after i.p. mCRP in the cortex (*p* = 0.006) and in the CA3 region (*p* = 0.02) only in ApoE4, but not ApoE2 and ApoE3, mice. n =11-14 in each group. 4C. Representative images of CD68^+^ active microglia (red), Iba1^+^ microglia (green) and GFAP^+^ astrocytes (purple) in the cortex of WT and ApoE knock-in mice after the i.p. treatment of PBS vs. mCRP are shown. Quantification of the microglial biomarkers CD68 (*p* = 0.02) and Iba1 (*p* = 0.0005) in the cortex showed significant differences between the PBS and mCRP groups only in ApoE4 mice, but the astrocyte biomarker GFAP (*p* = 0.12) only showed tendency in the ApoE4 mice. n =11-14 in each group. 4D. Double immunostaining of CD8 (red), CD3 (green), and double-positive cells (yellow) was performed to study the transcytosis of T lymphocytes in the cortex after i.p. treatment with PBS (left columns) vs. mCRP (right columns) in WT and different ApoE knock-in mice. Quantification of CD8^+^ T lymphocytes and CD8^+^/CD3^+^ T lymphocytes in the cortex and the comparisons between PBS vs. mCRP treatment in each genotype was conducted. mCRP significantly increased the number of T lymphocytes only in the WT (*p* = 0.05) and ApoE4 (*p* = 0.001) mice. n = 7-8 in each condition. 4E. Different correlation analyses using the data from all the mice used in the experiments were conducted to examine the relevance of different factors of the mCRP vs. ApoE to CD31 binding and the development of AD related pathology in the brain. The first row graphs show that the CD31 level (r = -0.48, *p* < 0.0001) was inversely related to PHF1 level in the brain, but PLA (ApoE-CD31) was not; the levels of PLA (mCRP-CD31) (r = 0.45, *p* = 0.02) and pCD31 (r = 0.40, *p* = 0.002) were positively correlated with PHF1 levels. The second row graphs show that the CD31 level (r = -0.35, *p* = 0.007) was inversely correlated with CD68/Iba1-positive cells in the brain, but PLA (ApoE-CD31) was not; the levels of PLA (mCRP-CD31) (r = 0.52, *p* = 0.02) and pCD31 (r = 0.45, *p* = 0.003) were positively correlated with double staining of CD68/Iba1^+^ cells. The third row graphs show that the number of CD8^+^/CD3^+^ T lymphocytes (y-axis) was negatively associated with CD31 level (r = -0.28, *p* = 0.03) and positively associated with the levels of pCD31 (r = 0.54, *p* < 0.0001); however, there was no association with PLA (ApoE-CD31) or PLA (mCRP-CD31). The forth row graphs show that the length of CD31^+^ microvessels (x-axis) was negatively correlated with the levels of pTau (r = -0.56, *p* < 0.0001), the numbers of CD68^+^/Iba1^+^ cells (r = -0.47, *p* = 0.0001), the numbers of CD8^+^/CD3^+^ T lymphocytes (r = -0.48, *p* < 0.0001) and tended to be correlated with the levels of Aβ1-42 (r = -0.21, *p* = 0.06) in the brain. Data are expressed as the mean ± SEM. One or two-way ANOVA with Tukey’s post hoc test and Pearson or Spearman correlation test were applied. \**p* < 0.05, \*\**p* < 0.01, \*\*\**p* < 0.001. The scale bar: 50 µm.

Since endothelia mediates migration of immune cells to tissues, we also examined and quantified levels of CD3^+^/CD8^+^ T lymphocytes in the cortex and hippocampus of ApoE mice. Figure 4D shows that elevated peripheral mCRP significantly increased the number of CD8^+^ and CD3^+^/CD8^+^ T lymphocytes in the cortex and hippocampus only in ApoE4 (p < 0.001) but not in ApoE3 and ApoE2 mice. In contrast, CD19^+^ B lymphocytes and CD14^+^ monocytes did not show significant changes after mCRP treatment across different ApoE genotypes (Supplement Figure 4B). Although CD31 is also expressed immune cells ^8^, unlike brain endothelia there was minimum or no colocalization of mCRP and CD3^+^/CD8^+^ T lymphocytes (Supplement Figure 4C), but the majority of CD3^+^ T immune cells were pCD31-positive cells (Supplement Figure 4D). These results mirror a recent study that demonstrated a large increased number of selective T lymphocytes found in the AD brain ^21^.

Upon collecting data from all the experimental mice (Figure 4E), tauopathy in the brain was significantly, positively correlated with mCRP-CD31 co-localization (r = 0.45, p = 0.02), but not linked with ApoE-CD31 co-localization. Consistently, tauopathy was negatively correlated with the CD31 expression levels (r = -0.48, p < 0.0001), but positively associated with the pCD31 levels (r = 0.40, p = 0.002) in the brain. In addition, the CD68^+^/Iba1^+^ microglia levels were significantly, positively with mCRP-CD31 co-localization (r = 0.52, p = 0.02) but not linked with ApoE-CD31 co-localization in the brain. Again, CD68^+^/Iba1^+^ microglia levels were negatively associated with CD31 expression (r = -0.35, p = 0.007) but positively with the pCD31 levels (r = 0.45, p = 0.0003) in the brain. Interestingly, brain CD3^+^/CD8^+^ T lymphocytes were not associated with either mCRP-CD31 or ApoE-CD31 binding. However, brain CD3^+^/CD8^+^ T lymphocytes were negatively correlated with CD31 expression (r = -0.28, p = 0.03), but positively correlated with pCD31 (r = 0.54, p < 0.001) in the brain. Furthermore, the average cerebrovascular length was negatively associated with tauopathy (r = -0.56, p < 0.0001), neuroinflammation marked by CD68^+^/Iba1^+^ microglia (r = -0.47, p = 0.0001), tended to be associated with Aβ1-42 (r = -0.21, p = 0.06) and significantly with CD3^+^/CD8^+^ T lymphocytes (r = -0.48, p < 0.001) in the brain. Importantly, the number of CD3^+^/CD8^+^ T lymphocytes was also associated with AD-associated pathological features (Supplement Figure 4E).

The above data suggest that mCRP-CD31 binding causes brain AD pathology and neuroinflammation through regulating CD31 expression and its phosphorylation (pCD31) leading to cerebrovascular damage with a severity pattern that mirrors the ApoE AD risk allele profiles (ApoE4>ApoE3>ApoE2).

### CD31 is a target of mCRP at the Alzheimer’s brain endothelia in humans

To further confirm the existence and the location of mCRP in the human brain, we used human temporal lobe tissues from 8 healthy controls and 10 AD patients and observed stronger mCRP immunostaining in the CD31^+^ cerebrovasculature of AD temporal cortex sections than in those of healthy control sections (Figure 5A). In contrast, ApoE levels in the CD31^+^ cerebrovasculature were significantly lower in AD brains compared with that in control brains (Figure 5B). Unlike cerebrovasculature ApoE, there was no difference in the ApoE levels in total brain tissues between AD and control brains (Supplementary Figure 5B). Further, using double staining and PLA, we found that AD brains tended to have higher levels of mCRP-CD31 binding (Figure 5A, Supplement Figure 5A) but lower levels of ApoE-CD31 binding than the control brains (Figure 5B), suggesting that CD31 is a competitive target for mCRP and ApoE in humans as well. Using western blots, we observed that there was a higher average level of pCD31 in AD cortex tissue than in control cortex tissue (Figure 5C) and that total CD31 levels tended to be lower in AD brains than in control brains, but this difference did not reach statistical significance (Figure 5A and 5C). Consistently with mouse data, the images of immunolabeled brain microvasculature show that pCD31 labeling in microvessels was significantly higher in AD brains than in control brains (Figure 5D). We noted that AD brains consistently exhibited microvessels with shorter lengths (Figure 5D) and more vessel termini (Supplement Figure 5B).

**Figure 5.**
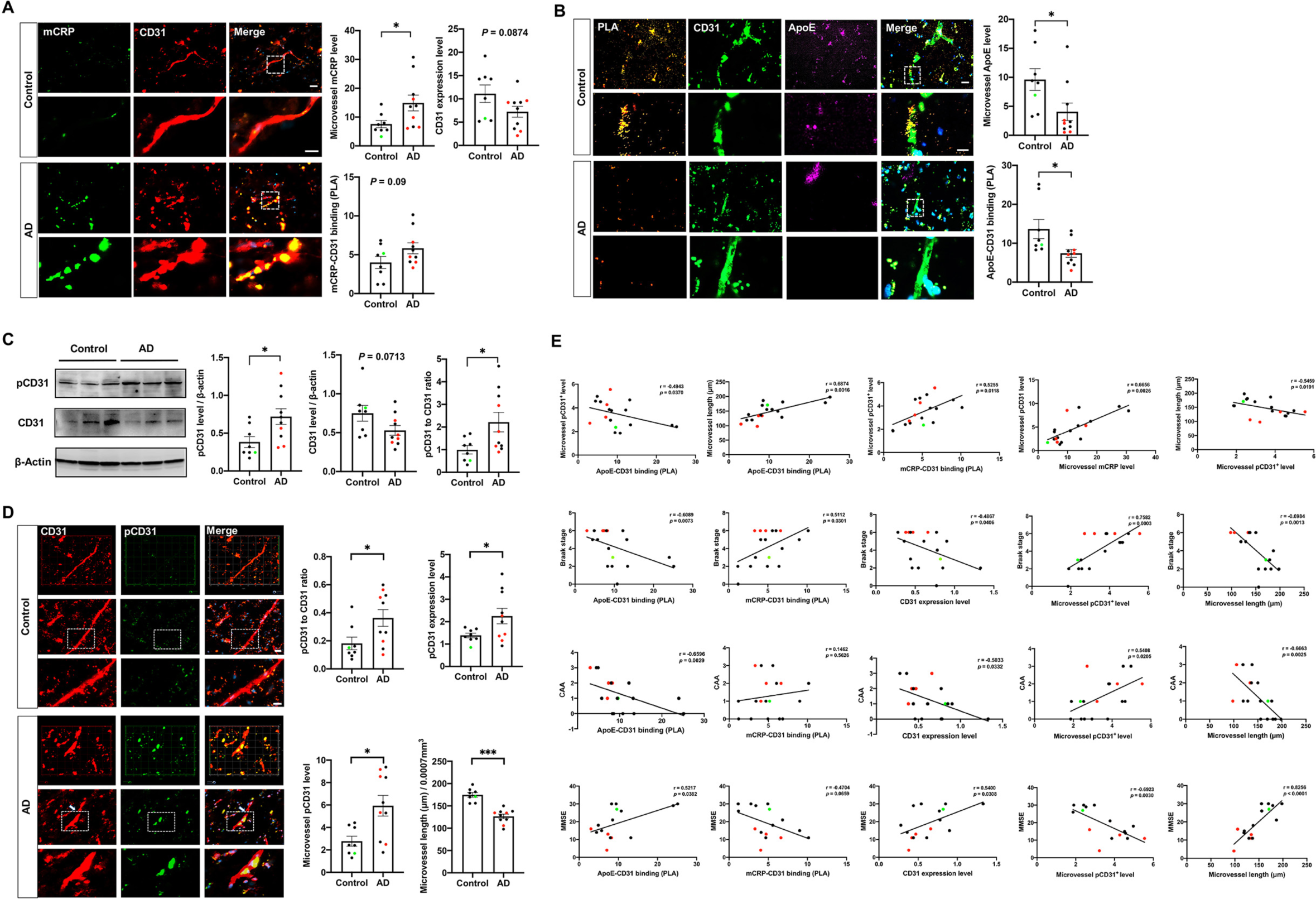
The relevant human data support the ApoE-CRP-CD31 pathway for AD pathogenesis in the brain. The human temporal cortex of healthy controls (n = 8) and AD patients (n = 10) was used for immunostaining. ApoE4 carriers (red dots) and ApoE2 carriers (green dots) are illustrated. 5A. Representative images of CD31^+^ microvessels (red), mCRP (green) and the merged images (yellow) in the temporal cortex are shown. The levels of microvessel mCRP and mCRP-CD31 binding were quantified and compared. AD brains had significantly higher mCRP immunostaining in microvessels (yellow) than control brains (*p*=0.01). AD brains tended to have lower levels of microvessel CD31 expression (*p* = 0.09) and higher levels of mCRP-CD31 binding detected by using PLA (*p* = 0.09) than control brains. 5B. PLA was also performed to detect interaction/binding between ApoE and CD31 in the temporal cortex of healthy controls and AD patients. Representative images of positive PLA (orange), ApoE (green) and CD31 (purple) staining are shown. The levels of microvessel ApoE and ApoE-CD31 were quantified and compared. AD brains had significantly lower levels of microvessel ApoE expression (*p* = 0.02) and ApoE-CD31 binding (*p* = 0.03) than control brains. 5C. Western blots for pCD31 and CD31 expression in the temporal cortex were performed and quantified after normalization against β-actin and compared between AD and controls. AD brains had higher levels of pCD31 (*p* = 0.02), tended to have lower levels of total CD31 (*p* = 0.07), and had a higher pCD31/CD31 ratio (*p* = 0.03) than control brains. 5D. Double immunostaining of temporal cortex sections with CD31 (red), pCD31 (green) and the merged images (yellow) were conducted to examine CD31 phosphorylation and microvessel integrity. AD brains had higher brain pCD31 levels (*p* = 0.05), pCD31/CD31 ratio (*p* = 0.03), and microvessel pCD31 (*p* = 0.01). Measurement of the lengths of CD31^+^ microvessels from 3D images revealed shorter lengths in AD brains than in controls (*p* < 0.001). 5E. Correlation analyses were conducted to investigate the mCRP-ApoE-CD31 pathway and brain AD pathology in humans. In the first row, the microvessel pCD31 level (y-axis) was positively associated with mCRP (r = 0.67, *p* = 0.003) and mCRP-CD31 binding (PLA) (r = 0.53, *p* = 0.01) and negatively associated with ApoE-CD31 binding (PLA) (r = -0.49, *p* = 0.04). Microvessel length (y-axis) was negatively associated with pCD31 (r = - 0.55, *p* = 0.02) but positively associated with ApoE-CD31 (r = 0.69, *p* = 0.002). In the second row, the Braak stage (y-axis) was negatively associated with CD31 expression (r = -0.49, *p* = 0.04), ApoE-CD31 binding (r = -0.61, *p* = 0.007) and microvessl length (r = - 0.70, *p* = 0.001); in contrast, the Braak stage was positively associated with mCRP-CD31 binding (r = 0.51, *p* = 0.03) and microvessel pCD31 expression (r = 0.76, *p* = 0.0003). In the third row, the CAA (y-axis) was negatively associated with CD31 expression (r = - 0.50, *p* = 0.03), ApoE-CD31 binding (r = -0.66, *p* = 0.003) and microvessl length (r = - 0.67, *p* = 0.003); in contrast, CAA was not associated with the level of mCRP-CD31 binding but was positively associated with microvessel pCD31 expression (r = 0.54, *p* = 0.02). In the fourth row, the Mini-Mental State Exam (MMSE) score (y-axis) was positively associated with CD31 expression (r = 0.54, *p* = 0.03), the level of ApoE-CD31 binding (r = 0.52, *p* = 0.04) and microvessl length (r = 0.83, *p* < 0.0001); in contrast, MMSE score tended to be negatively associated with mCRP-CD31 binding (r = -0.47, *p* = 0.07) and significantly associated with the level of microvessel pCD31^+^ cells (r = -0.69, *p* = 0.003). Data are expressed as the mean ± SEM. Student’s t-test and Pearson or Spearman correlation tests were used. \**p* < 0.05, \*\**p* < 0.01, \*\*\**p* < 0.001. The scale bars are 100 µm, 20 µm, and 10 µm.

To verify potential causal relationships between the ApoE, mCRP and CD31 in cerebrovasculature and the neuropathological and clinical phenotypes of AD, we performed correlation analyses (Figure 5E). The microvessel pCD31 level in the brain was negatively correlated with ApoE-CD31 binding (r = -0.46, p = 0.04) but positively correlated with both the microvessel mCRP level (r = 0.66, p = 0.003) and mCRP-CD31 binding (r = 0.53, p = 0.01) in the brain. The microvessel lengths in the brain was positively association with ApoE-CD31 (r = 0.69, p = 0.002) but negatively association with the level of pCD31 (r = -0.55, p = 0.02) in the brain. Furthermore, the Braak stage of AD pathology was negatively associated with the CD31 expression level (r = -0.49, p = 0.04), ApoE-CD31 binding (r = -0.61, p = 0.007) and microvessel lengths (r= -0.70, p = 0.001), whereas positively with mCRP-CD31 binding (r = 0.51, p 0.03) and pCD31 level (r = 0.76, p = 0.0003). Cerebral amyloid angiopathy (CAA) was negatively associated with the CD31 expression level (r = -0.50, p = 0.03), ApoE-CD31 binding (r = -0.66, p = 0.003) and microvessel lengths (r = -0.67, p = 0.003), whereas positively associated with mCRP-CD31 binding and pCD31 levels (r = 0.54, p = 0.02). Consistently with the pathological traits observed in the brains, the MMSE scores when the patients were alive were positively associated with the CD31 expression level (r = 0.54, p = 0.03), ApoE-CD31 binding (r = 0.52, p = 0.04) and microvessel lengths (r = 0.83, p < 0.001), but negatively associated with mCRP-CD31 binding (r = -0.47, p = 0.07) and pCD31 levels (r = -0.69, p = 0.003). These results demonstrate a significant, and inverse relationship between mCRP-CD31 binding and ApoE-CD31 binding on AD pathology, with mCRP-CD31 increasing phosphorylation of CD31 and the degree of cerebrovascular pathology in the brains of AD patients, and ApoE-CD31 having the opposite effect in human ApoE2 (green dots) vs. ApoE4 (red dots) carriers.

### mCRP expression results in neuroprotective or neurodegenerative pathway activation in an ApoE-dependent manner

To understand the molecular pathways affected by mCRP in an ApoE-specific manner, we used flow cytometry to isolate CD31^+^ BECs from three ApoE2, ApoE3 and ApoE4 knock-in mice after i.p injection of PBS or mCRP for six weeks (Figure 6A schematic). We then used RNA sequence analysis to identify the *in vivo* prominent factors and important molecular pathways affected mCRP within distinct ApoE genotypes. Here, all the genes were ranked by the Wald-test statistic computed between mCRP and PBS for each ApoE genotype, followed by GSEA analysis. We identified sets of genes that are coordinately up- and down-regulated with respect to expression changes between mCRP and PBS in each ApoE genotype group. The most highly enriched gene sets upregulated in the cerebral endothelia of ApoE4 mice after mCRP expression reflected changes in pathways including 1) oxidative phosphorylation, 2) mitochondria metabolism/respiration, 3) mTORC1 signaling, and 4) Alzheimer’s disease; in contrast, these pathways were downregulated by mCRP in the endothelia of ApoE2 brains (Figure 6B). On the other hand, mCRP significantly upregulated other pathways in the endothelia of ApoE2 brain, including 1) epigenetic histone methylation, 2) synapse formation, 3) Notch signaling and 4) vasculogenesis; in contrast, mCRP downregulated these pathways in the endothelia of ApoE4 brains (Figure 6C). The enrichment scores of these pathways in the ApoE2 group were inversely correlated with those in ApoE4, while the ApoE3 comparisons yielded intermediate enrichment scores (Supplementary Figure 6A).

**Figure 6.**
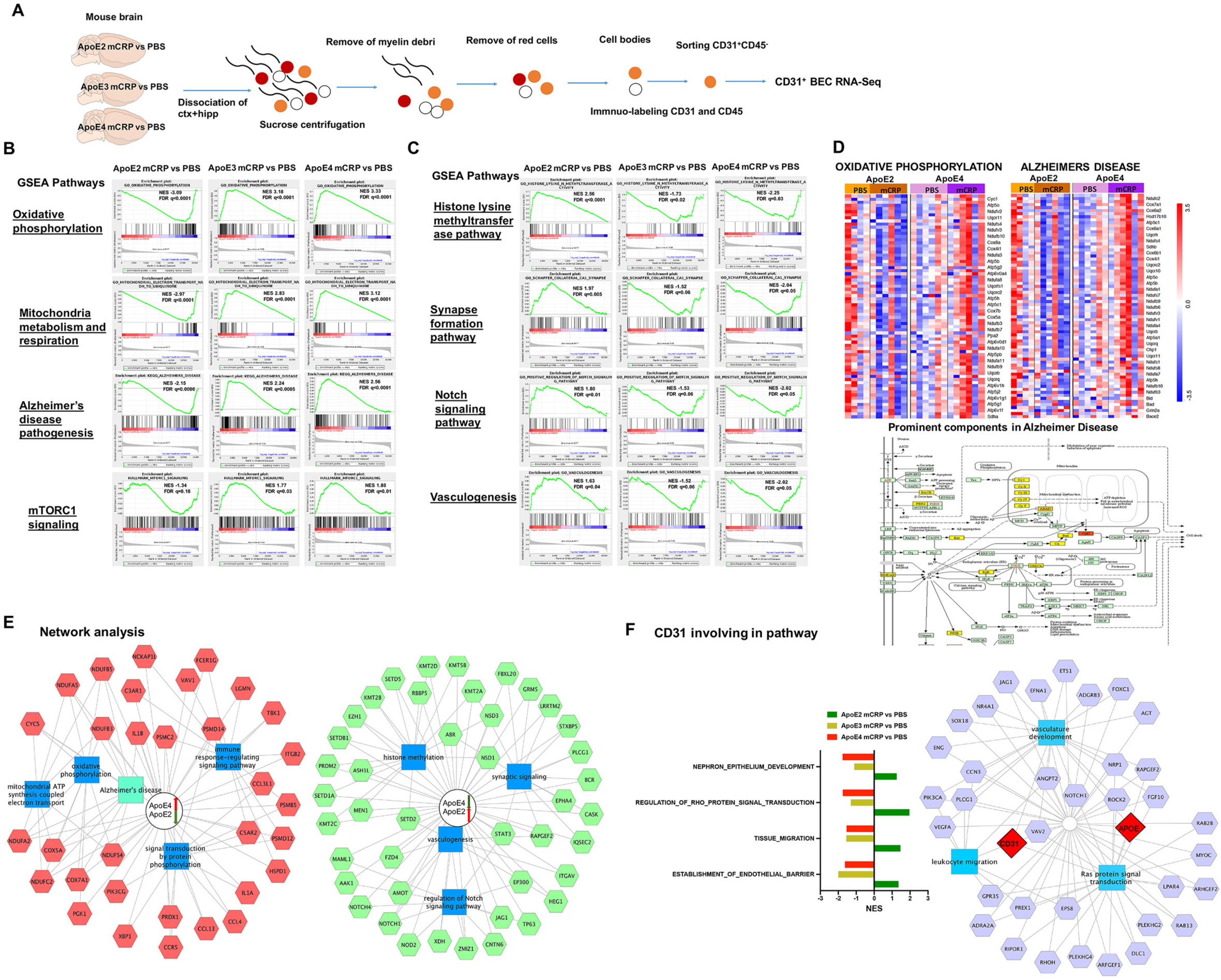
mCRP impacts the brain endothelial cell pathways in an ApoE genotype-dependent manner. 6A. The schematic showed CD31^+^ brain endothelial cells (BECs) were isolated by flow sorting and subjected to RNA sequencing from the mouse brain treated with i.p. PBS vs. mCRP. 6B-D. GSEA based on their differential gene expression was conducted to study the prominent pathways enriched in gene lists from different ApoE genotypes for AD pathogenesis. For the comparison between mCRP and PBS in each genotype, genes were ranked by GSEA based on their differential expression level. Four pathways, including 1) oxidative phosphorylation, 2) mitochondrial metabolism and respiration, 3) Alzheimer’s disease and 4) mTORC1 signaling, were upregulated by mCRP in ApoE4 mice but were downregulated in ApoE2 mice, while ApoE3 mice were in the middle (6B). In contrast, another four pathways, including 1) the histone lysine methyltransferase pathway, 2) synapse formation, 3) Notch signaling and 4) vasculogenesis, were upregulated by mCRP in ApoE2 mice but downregulated in ApoE4 mice, while ApoE3 mice were in the middle (6C.) The top or bottom of the ranked gene list in each ApoE comparison was evaluated using the enrichment score (green line) to determine whether a candidate pathway was significant. Black vertical lines mark positions where members of a particular pathway appear in the ranked list of genes. NES, normalized enrichment score; FDR, false discovery rate and q values. 6D shows the heatmaps of z-scored gene intensities of the differentially expressed genes enriched in the pathways/modules of oxidative phosphorylation and Alzheimer’s disease for ApoE2 and ApoE4 CD31^+^ BECs after the i.p. treatment. The key factors involved in the Alzheimer’s disease pathway that overlapped with the oxidative phosphorylation pathway were ranked and labeled by different intensities of yellow color. 6E. The RNAseq data were used for the functional enrichment analysis of the different signatures between ApoE4 and ApoE2 after the i.p. treatment of PBS vs. mCRP by using the ToppCluster tool (FDR correction, *p* < 0.05). The red/green box represents the top genes with up/downregulated expression in the ApoE4 mCRP vs PBS comparisons and down/upregulated expression in the ApoE2 mCRP vs PBS comparisons. The genes were connected to form a hub biological process and pathway (blue box). 6F. The CD31-related pathways are listed and shown according to NES comparison between mCRP vs PBS in each ApoE genotype (left panel). The network of CD31 and ApoE together with other components involved in biological processes, including vasculogenesis, leukocyte migration and Ras signaling pathways, is illustrated (right panel). Data are expressed as the mean ± SEM. One or two-way ANOVA with Tukey’s post hoc test was used. \**p* < 0.05, \*\**p* < 0.01, \*\*\**p* < 0.001; ^#^*p* < 0.05, ^##^*p* < 0.01, ^###^*p* < 0.001.

Figure 6D shows the heatmaps of the top differentially expressed genes in the pathways of oxidative phosphorylation and AD in the CD31^+^ BEC of ApoE2 and ApoE4 mice treated with PBS vs. mCRP. Most of these genes such as CYTC, which are involved in the oxidative phosphorylation pathway in mitochondria and also play a key role in AD pathogenesis, showed increased expression in ApoE4 mice treated with mCRP but reduced expression in ApoE2 mice treated with mCRP.

We further performed functional network analysis according to the ranked gene list (Figure 6E and Supplement Figure 6B). ApoE4-specific signature genes are featured prominently in the processes of oxidative phosphorylation (CYCS, COX5A/7A), mitochondrial ATP synthesis (NDUFA and NDUFB, NDUFC), the immune response (IL1A, IL1B, CCL3L1) and protein phosphorylation (PRDX1, PIK3CG). ApoE2-specific signature genes functioned mainly in histone methylation (NSD1, SETD2), synaptic signaling (KMT2A, STAT3), Notch signaling (NOTCH1, EP300) and vasculogenesis (ZMIZ1, XDH). We extracted CD31-associated gene ontologies and found that CD31-associated signaling pathways were upregulated in ApoE2 but downregulated in ApoE4 and ApoE3. Network analysis illustrates that CD31, ApoE and other candidate components were mainly involved in the biological processes of vascular development, leukocyte migration and Ras signal transduction (Figure 6F). Taken together, these data suggest that when peripheral inflammation occurs, ApoE2 exerts protective effects by enhancing vascular protection and repair, whereas ApoE4 exerts deleterious effects by disrupting mitochondrial function and vascular inflammation, which leads to cerebrovascular damage and AD pathogenesis.

## Discussion

This study demonstrates a novel pathological pathway, the ApoE-mCRP-CD31 pathway, in the blood-facing endothelia of the BBB that differentially affects cerebrovasculature and AD pathogenesis in an ApoE allele-specific fashion, and that is likely exacerbated when peripheral chronic inflammation occurs (Figure 7 schematic). We find that mCRP binds to endothelial CD31, competing with ApoE alleles for CD31 which had ApoE4 equivalent to ApoE knockout (Figures 1 and 2). Elevated peripheral mCRP leads to decreased CD31 expression and increased phosphorylation of CD31 (pCD31), caused cerebrovasculature damage marked by shortened microvessel lengths (Figure 3), leading to the extravasation of T lymphocytes and the development of AD pathogenesis particularly in the ApoE4 brain (Figure 4). This process was antagonized by cerebrovascular ApoE-CD31 binding most effectively in the ApoE2 brain (Figures 1-2 and 5-6).

**Figure 7.**
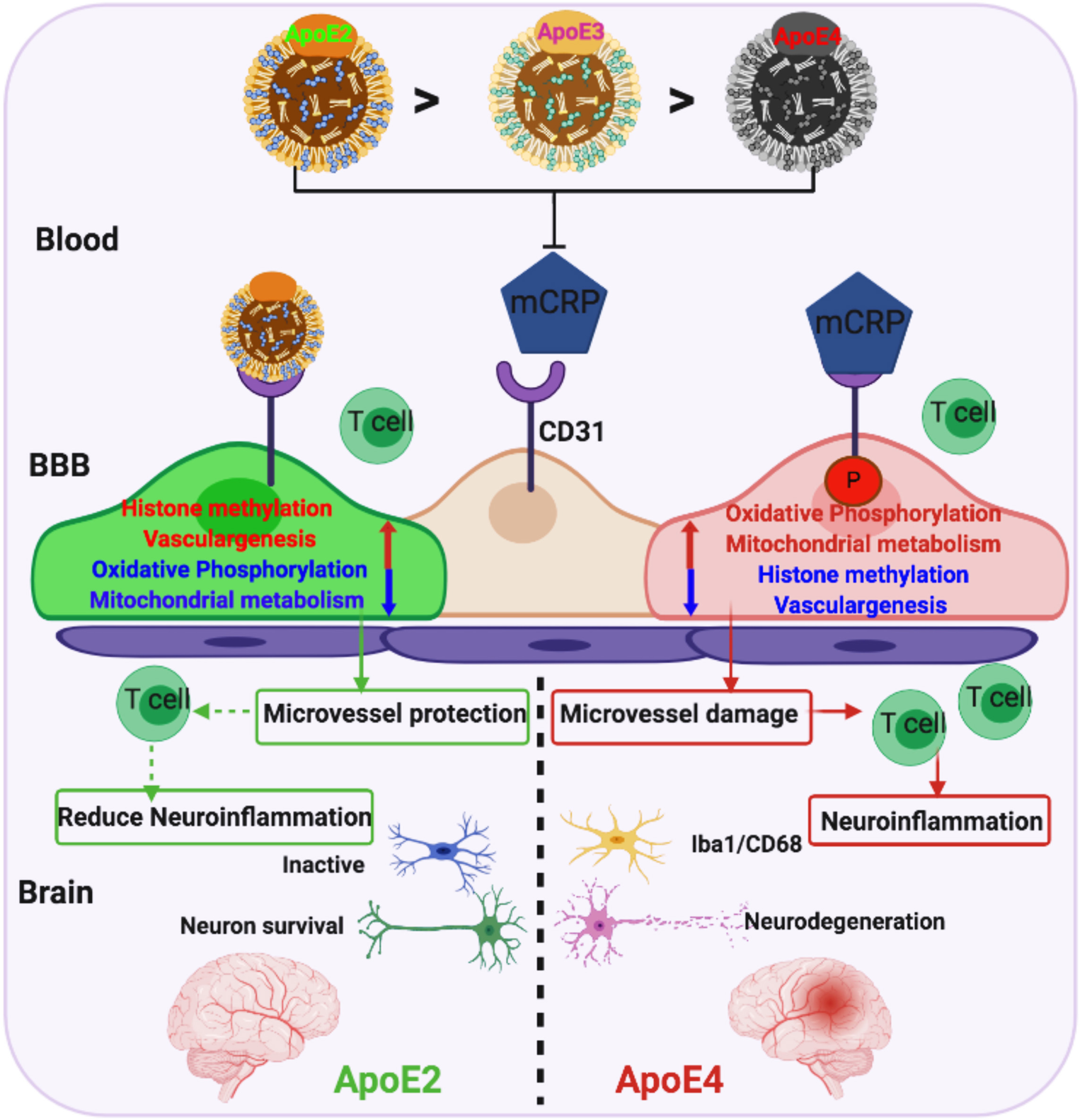
A model illustrating the differential responses of ApoE2 vs. ApoE4 carriers to mCRP and the differential regulation of mCRP-induced cerebrovascular neuroinflammation leading to AD pathogenesis in the brain. This study demonstrated a novel pathological mechanism, the competition of ApoE and mCRP to CD31binding, for cerebrovascular neuroinflammation resulting in an early stage of AD pathogenesis in the brain. During the chronic stage of peripheral inflammation, pCRP proteins disassociate into mCRP. mCRP binds to CD31 on microvessels to increase CD31 phosphorylation (pCD31), cause damage to the cerebrovasculature and induce extravasation of T lymphocytes into the brain, leading to AD pathogenesis (ApoE4>ApoE3>ApoE2). This process is antagonized by ApoE-CD31 binding (ApoE2>ApoE3>ApoE4) to block mCRP-CD31 binding and differentially regulate pathways (mitochondrial function, epigenetics and vasculogenesis) to intervene in the neurodegenerative process of AD.

Peripherally elevated mCRP caused the highest mCRP deposits in the ApoE4 brains compared to ApoE3 and ApoE2 brains, probably leading to a lower blood CRP level in ApoE4 carriers (Supplementary Figure 1). While these mice do not have transgenic genes for either tau or Aβ, in response to peripheral mCRP, our study illustrated an ApoE4-related AD pathway at an early stage through peripheral chronic inflammation compared to ApoE2 and ApoE3 mice (Figure 4). In parallel, human AD brains have a high level of mCRP-CD31 binding in the cerebrovasculature (Figure 5A). Consistently, other studies have shown that mCRP colocalizes with amyloid pathology in the human AD brain ^5, 22^. While ApoE4 has been shown to have enhancing effects to induce neuroinflammation ^23^, chronic peripheral inflammation induced cytotoxic effects and increased the occurrences of neuroinflammation and the severity of neurodegeneration ^24^. Additionally, CRP levels are known to increase with age ^4^, and mCRP has been shown to play a role in the pathogenesis of peripheral vascular diseases including cardiovascular diseases ^25^ and post-stroke inflammation ^26^.

Our study demonstrates the cerebrovascular changes during peripheral chronic inflammation through the ApoE-mCRP-CD31 interactions. ApoE proteins and mCRP competitively bound to endothelial CD31 and that there was the lowest level of ApoE-CD31 binding in the presence of ApoE4 leading to more mCRP binding to CD31 on blood-facing endothelial cells in the brain (Figures 1, 2 and 5). mCRP increased pCD31 production, and this was further linked with shortened CD31^+^ cerebrovasculature predominantly in the ApoE4 brain, and less so in ApoE3 and ApoE2 brains (Figures 3 and 5). mCRP-CD31 binding and pCD31 levels were also found to be linked with CAA in humans (Figure 5E), the major cerebrovascular pathology in human AD ^27^. As mCRP induced neuroinflammation (Figure 4C) in ApoE4 mice, phosphorylation of the CD31 cytoplasmic domain ^28^ has been shown to limit the inhibitory response of CD31 to inflammatory reactions ^8, 19^. Consistently, elevated mCRP caused the changes of several cerebrovascular biomarkers. The levels of vWF, a factor released from damaged vasculature ^20^, were negatively correlated with vascular length and significantly increased by mCRP selectively in the ApoE4 brain (Figure 3B). eNOS, especially p-eNOS, is a protective factor in the vasculature and dementia by being responsible for most of the vascular nitric oxide produced ^29^ ^30^. Interestingly, mCRP induced the p-eNOS level particularly in ApoE4-expressing cells and further increased the p-eNOS level in CD31 knockdown endothelia (Figure 3D), suggesting that p-eNOS is a downstream compensating factor, but NF-κB and vWF are the downstream deteriorating factors, in the ApoE-mCRP-CD31 pathway for cerebrovascular inflammation. Consistently, another study shows that ApoE4-positive endothelial cells derived from human stem cells have higher levels of Aβ, vWF and cytokines than ApoE3 endothelial cells, suggesting that the ApoE isoforms affects endothelial cell function ^31^, while the ApoE4 allele also disrupts brain-facing cells, e.g pericytes and astrocytes, of the BBB ^32, 33^. CD31^+^ endothelial cells are also shown to be vascular stem cells ^34^ and probably are the precursors of pericytes for BBB ^35^.

Our study discovered that peripheral mCRP increased the number of T lymphocytes and possibly monocytes in the ApoE4 brains (Figure 4D). This finding is supported by a recent paper demonstrating that the number of clonally expanded CD8 T cells is high in the cerebrospinal fluid of AD patients ^21^. Although CD31 is expressed in both endothelial and immune cells in peripheral tissues ^36, 37^ ^38, 39^, mCRP was found to bind endothelial CD31 in ApoE4 brains (Figure 1) but not to CD31 in migrated T lymphocytes in the brain (Supplement Figure 4). Since CD31 expressed in peripheral endothelial cells plays roles in regulating endothelial peripheral vascular junction/integrity ^8, 19^, our study suggests that mCRP modulates extravasation of T lymphocytes mainly through binding to brain endothelial CD31 based on the following results: 1) mCRP significantly increased CD31 phosphorylation and decreased the expression of CD31 in primary endothelia (Figure 1). Lack of CD31 is associated with increased mononuclear leukocyte translocation into the CNS in a mouse model of experimental autoimmune encephalomyelitis (EAE) ^40^. 2) CD31 protein expression is reduced in the AD brain (Figure 5). Endothelial culture from AD patients increase monocyte transmigration compared to those from normal controls, and Aβ treatment enhances this transmigration ^41^, as microvascular endothelia contraction can be accompanied by leukocyte extravasation in peripheral tissues ^42^.

It is intriguing that the molecular pathways triggered by mCRP-CD31 binding demonstrated oppositional regulation in ApoE4 vs. ApoE2 endothelia (Figure 6). Although blood-facing endothelia in the brain express little ApoE, endothelia can be exposed to, and influenced more by, ApoE protein in blood, and human studies show that lower plasma concentrations of ApoE are associated with brain AD pathology ^43^.

Notably, after mCRP stimulation *in vivo*, ApoE4 BECs showed significantly increased, but ApoE2 BECS decreased, gene expression related to four pathways linked to neurodegenerative processes 1) oxidative phosphorylation, 2) mitochondrial metabolism, 3) mTORC1 signaling and 4) AD pathology (Figure 6B and D-E). On the other hand, ApoE2 BECs showed upregulation, but ApoE4 BECs showed downregulation, of four other pathways more commonly associated with neuroprotection 1) histone methylation-epigenetics, 2) synapse formation, 3) Notch signaling and 4) vasculogenesis (Figure 6C and E). Interestingly, Rho signaling was linked with both CD31 and ApoE in endothelial network modules of vascular development and leukocyte migration by mCRP treatment (Figure 6F). Deficient regulation of Rho signaling contributes to impaired endothelial barrier integrity and elevated p-eNOS levels and that Ras/ERK signaling plays an important role in T cell migration, which is regulated by eNOS ^37^. RhoA/Rho kinase (ROCK) is also involved in eNOS function ^44^, as its activation decreases eNOS expression to disrupt endothelial barriers ^45^. Signals mediated by CD31 in the endothelium are both necessary and sufficient to prevent endothelial cell death and confer immune privilege to protect the vascular endothelium after inflammatory attacks ^46^. These data are consistent with several published results on AD pathological pathways. Oxidative phosphorylation provides an energy-generating pathway relying on mitochondria for ATP production to maintain cerebral vasculature ^47^. Mitochondrial dysfunction in endothelial cells has been implicated in mediating BBB failure in degenerative disorders, such as AD ^48^. A study indicated that ApoE deficiency plus adverse factors could alter histone lysine methylation modifications in vascular endothelial cells ^49^. It has been shown that Notch pathway genes, such as Notch1 and Jag1, are downregulated during the response to vascular injury ^50^. While mCRP activates Notch to promote angiogenesis through PI3K ^51^, in contrast, after Notch signaling deactivation, endothelial cells lose their barrier function, which leads to endothelial hyperpermeability, leakage and inflammatory responses ^52^.

In summary, and as illustrated in Figure 7, our study provides evidence for an ApoE-mCRP-CD31 pathway in blood-facing endothelia of the BBB that can regulate responses to mCRP in peripheral chronic low-grade inflammation and impact AD pathology. Through different abilities to compete with mCRP for CD31, ApoE2 appears to protect the cerebrovasculature more potently than ApoE4. Combined, our data suggest that ApoE2 protects via a mechanism that involves upregulating epigenetic modifications, Notch signaling and vasculogenesis. In contrast, ApoE4 exhibits effects on the cerebrovasculature that are more detrimental than ApoE3 and ApoE2. In the ApoE4 background equivalent to ApoE knockout, mCRP increases CD31 phosphorylation to promote cerebrovascular damage via disrupting mitochondrial metabolism, enhancing oxidative phosphorylation and increasing the immune response, leading to cerebrovascular neuroinflammation and AD pathology. Given the high frequency at which elderly people experience peripheral inflammatory attacks and develop chronic low-grade inflammation, which results in the formation and release of mCRP, these data may explain why some, but not all, ApoE4 carriers develop AD by the age of 90, and why ApoE2 is protective against the disease ^4, 53^.

## ACKNOWLEDGEMENTS

This work was supported by grants from NIA, R21AG045757, RO1AG-022476, R01 AG054546 and R56AG059805 (W.Q.Q). Hua Tian was supported by Chinese National Foundation for Abroad Scholarship.

We are grateful to Dr. Adam Gower for his RNAseq data analyses and advice on the manuscript writing. We thank staff from Boston University School of Medicine Microarray and Sequencing Center for their RNA sequencing and professionism. We also want to express our thanks to the FHS and Boston University Alzheimer’s Disease Center (BUADC) participants for their decades of dedication and to the staff for their hard work in collecting and preparing the data and brain tissues. FHS was supported by the National Heart, Lung, and Blood Institute contract (N01-HC-25195) and by grants from the National Institute of Neurological Disorders and Stroke (NS-17950) and from the National Institute on Aging (AG-008122, AG-16495, AG-022476). Support for BU ADC was provided through NIA P30 AG13864.

## Contributors

Z.Z. and W.Q. designed the research; Z.Z., H.N. and Q.G. performed the research; Z.Y., J.Y., S.Z., H.T. and Q.L. contributed to data analysis; I.R., B.W. and L.P. provided the reagents; Y.A. contributed to RNA sequencing; J.B., B.W. and A. E. contributed to project suggestion; T. S. provide the human samples. J.H., Q.T. and X.Z. analyzed the human data; Z.Z. and W.Q. were responsible for results discussion as well as manuscript preparation.

## Supplementary Figure legends

**Supplement Figure 1. mCRP alters CRP, endothelial CD31 and pCD31 in the ApoE4 brain.**

S1A. The levels of CRP in human serum samples (left panel) and mouse serum samples (right panel) based on wild-type (WT) vs. different ApoE genotypes are shown. The comparisons with statistical significance are indicated.

S1B. Intraperitoneal (i.p.) injections of purified human CRP, which is mainly composed of pCRP and a small amount of mCRP, or recombinant human mCRP into WT mice and mice expressing different ApoE genotypes. Representative images of the cortex with immunofluorescence staining by anti-pCRP and anti-mCRP antibodies are shown.

S1C. Different mice were treated with i.p. injection of recombinant mCRP three days per week for 6 weeks. Serum mCRP levels were detected by ELISA (left panel); brain mCRP deposits were measured by the quantitation of immunofluorescence staining (middle panel, *p =* 0.05). There was a negative correlation between blood and brain mCRP levels in all the experimental mice (right panel, *p =* 0.03). n = 11-14 mice in each group.

S1D. Mouse CRP levels in the hippocampus were measured by ELISA. The left graph shows the quantitation of brain CRP levels across mice with different ApoE genotypes in the control i.p. PBS groups. (WT/ApoE2/ApoE3 vs. ApoE4, *p* = 0.04, *p* = 0.03, *p* = 0.04). The right graph shows the fold changes of brain CRP after the i.p. mCRP treatment normalized by the average level from the PBS treatment in each genotype. (WT/ApoE2/ApoE2 vs. ApoE4, *p* = 0.02, *p* = 0.006, *p* = 0.01). n = 9-10 in each group.

ApoE4 mice had the lowest level of CRP but higher fold changes of CRP after the i.p. mCRP treatment.

S1E. The total CD31 and pCD31 levels were quantified by western blots and normalized against β-actin, and the ratio of pCD31 to CD31 was calculated. There were significant changes in CD31 and the pCD31/CD31 ratio only in ApoE4 mice after mCRP treatment. S1F. Different concentrations of mCRP were added to primary CD31^+^ BECs for a 24-hour incubation followed by triple immunostaining with mCRP (purple), CD31 (green) and pCD31 (red) antibodies. The nuclei were stained with DAPI. The scale bar is 50 µm. Data are expressed as the mean ± SEM. One or two-way ANOVA with Tukey’s post hoc test, two-sided; \**p* < 0.05, \*\**p* < 0.01, \*\*\**p* < 0.001.

**Supplement Figure 2. The characterizations of their interaction between mCRP, ApoE**

S2A. Proximity ligation assays (PLAs) were performed. No binding/colocalization of mCRP and ApoE was detected on the cortex sections from mice with different ApoE genotypes regardless of i.p. mCRP treatment. The nuclei were stained with DAPI (blue). The scale bar is 50 µm.

**Supplement Figure 3. The mCRP deposits and CD31+ microvessels on hippocampal sections of different genotype mice after elevating peripheral mCRP.**

Representative images of hippocampal areas CA1 (S3A), CA3 (S3B) and DG (S3C) stained with CD31 (red) and mCRP (green) and merged (yellow) are shown to observe mCRP deposits and microvessel integrity. The quantifications of mCRP (green color) and CD31 (red color) fluorescence intensities were compared among different genotype mice with/without i.p. mCRP treatment (S3D).

Data are expressed as the mean ± SEM. One or two-way ANOVA with Tukey’s post hoc test were used. \**p* < 0.05, \*\**p* < 0.01, \*\*\**p* < 0.001. The scale bar is 50 µm.

**Supplement Figure 4 Peripheral mCRP’s effects on amyloid β expression level and immune cells in the brain across ApoE genotypes.**

S4A. Mouse amyloid β (Aβ)1-42 levels in the hippocampus and the cortex were measured by ELISA. The fold changes of Aβ42 were normalized by using the average level measured following the PBS treatment in each genotype (WT, ApoE2, ApoE3 and ApoE4, n = 9-10 in each genotype). Only the hippocampal region showed a slight increase in Aβ1-42 levels in ApoE4 mice compared with ApoE2 mice after mCRP treatment (*p* = 0.003).

S4B.Immunostaining of cortex sections for CD19 (red, a B cell marker) and CD14 (green, a monocyte marker) positive cells are shown from WT and ApoE knock-in mice treated with PBS (left columns) and mCRP (right columns). The fluorescence intensity of CD19^+^ and CD14^+^ cells in the cortex was quantified, and the comparisons between PBS and mCRP treatments did not reach statistical significance in all genotypes. n = 7-8 in each condition. S4C. Triple immunostaining of mCRP (purple) and T-lymphocyte marker CD8 (red) and CD3 (green) was conducted in the cortex. Correlation coefficients and R values were calculated and shown.

S4D. Double immunostaining of pCD31 (green) and CD3 (red) in the cortex and the CA3, DG and CA1 areas of the hippocampus were conducted. The overlapping signals indicate that pCD31 expression also colocalized with immune T lymphocytes marked by CD3.

S4E. The graphs show that the number of CD8^+^/CD3^+^ T lymphocytes (x-axis) was positively correlated with the levels of pTau (r = 0.47, *p* = 0.0001), the levels of Aβ1-42 in the hippocampus (r = 0.47, *p* = 0.006) and the number of CD68^+^/Iba1^+^ cells (r = 0.53, *p* < 0.0001) in the brain.

Data are expressed as the mean ± SEM. One or two-way ANOVA with Tukey’s post hoc test and Pearson or Spearman correlation test were used. \**p* < 0.05, \*\**p* < 0.01, \*\*\**p* < 0.001. The scale bar is 50 µm.

**Supplement Figure 5. Characterization of pathological indicators of the mCRP-ApoE-CD31 pathway in human brains.**

S5A. PLA was conducted to detect interaction/binding between mCRP and CD31 on cortex sections from healthy controls and AD patients. Representative images of positive PLA (orange), mCRP (purple) and CD31 (green) staining are shown. All the mCRP signals (purple) on the cortex section, including microvessels and brain tissues, were quantified and compared between the control and AD groups (*p* = 0.12).

S5B. The total ApoE signals (purple) on the cortex section were quantified and compared between the control and AD groups (*p* = 0.22). The numbers of pCD31^+^ cells (pCD31^+^ level) in microvessels were quantified and found to be significantly increased in AD brains compared with control brains (*p* = 0.0002). AD brains were observed to have more vessel terminals than control brains (*p* = 0.04).

Data are expressed as the mean ± SEM; the data from the *APOE4* allele are labeled in red, and the *APOE2* allele data are labeled in green; Student’s t-test; \*\*\**p* < 0.001. The scale bars are 20 µm and 10 µm

**Supplement Figure 6. mCRP effects on brain endothelial cell pathways in an ApoE genotype-dependent manner.**

S6A. Different ApoE knock-in mice were treated with i.p. PBS vs. mCRP for 6 weeks followed by isolation of brain CD31^+^ BECs and RNAseq. Correlation analyses were conducted to examine the relationship between ApoE4 and ApoE2/ApoE3 CD31^+^ BECs in NES. There was a reverse correlation between the ENS of ApoE4 and ApoE2 and a positive correlation between the ENS of ApoE4 and ApoE3. This result indicated that after mCRP treatment, the differential gene enrichment pathways in ApoE4 BECs were significantly regulated in an opposite manner compared with those in ApoE2 BECs, whereas they were regulated in parallel with those in ApoE3 BECs. Pearson correlation tests were conducted.

S6B. Differentially regulated pathways between the ApoE4 and ApoE2 comparisons are shown, including top pathways, energy metabolism and chromosome organization and pathological pathways, leukocyte immunity and endothelial vasculature. The NES of ApoE3 is mostly in the middle.

