## Supplemental Figures for "Competition between distinct ApoE alleles and mCRP for the endothelial receptor CD31 differentially regulates neurovascular inflammation and Alzheimer’s disease pathology"

**Supplemental Figure 1**

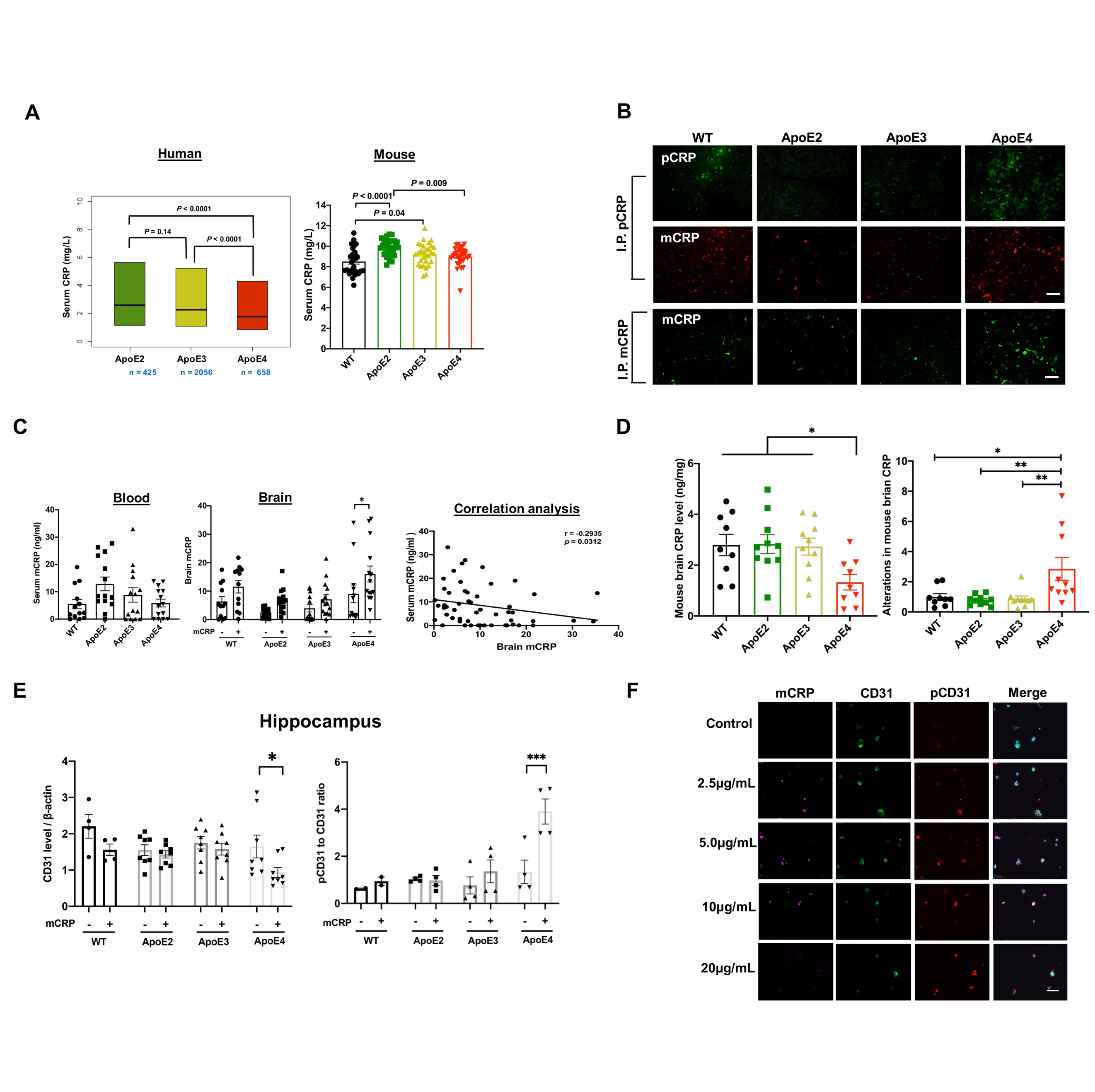

**Supplemental Figure 2**

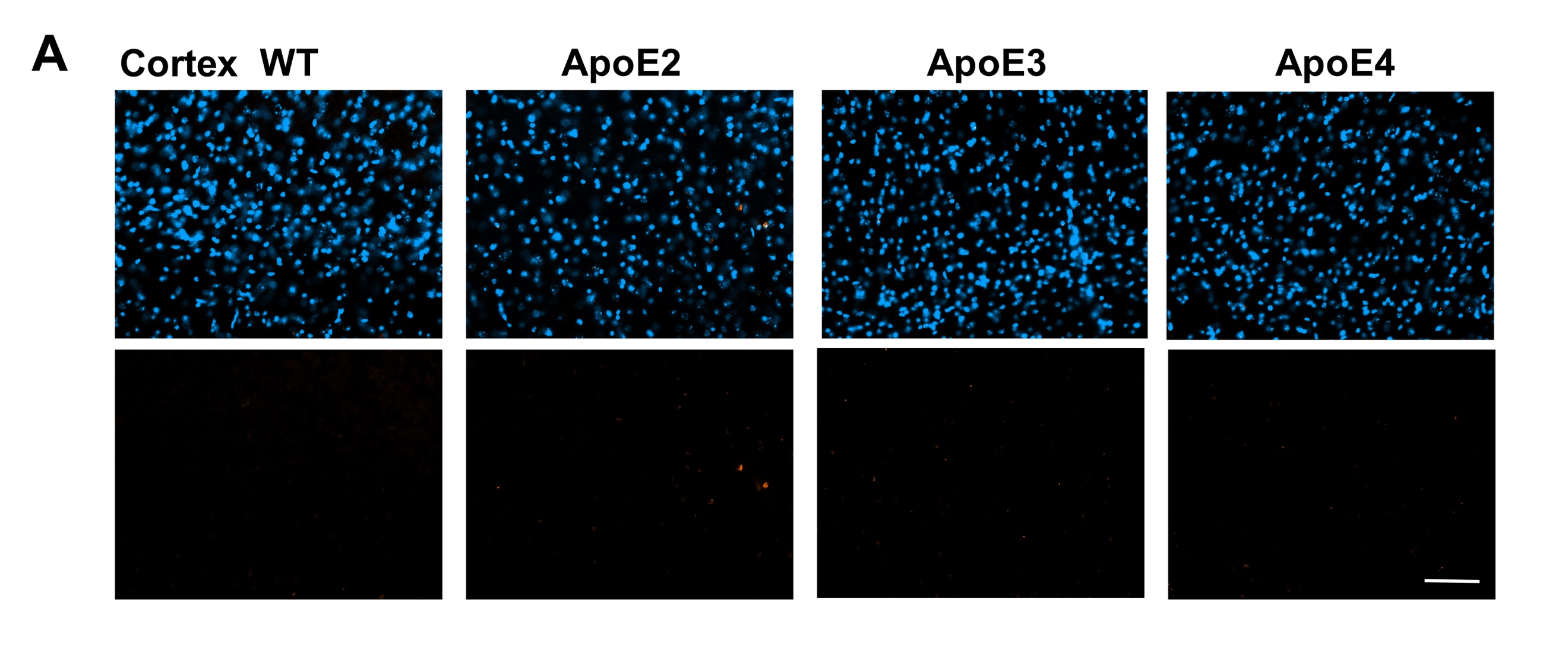

**Supplemental Figure 3**

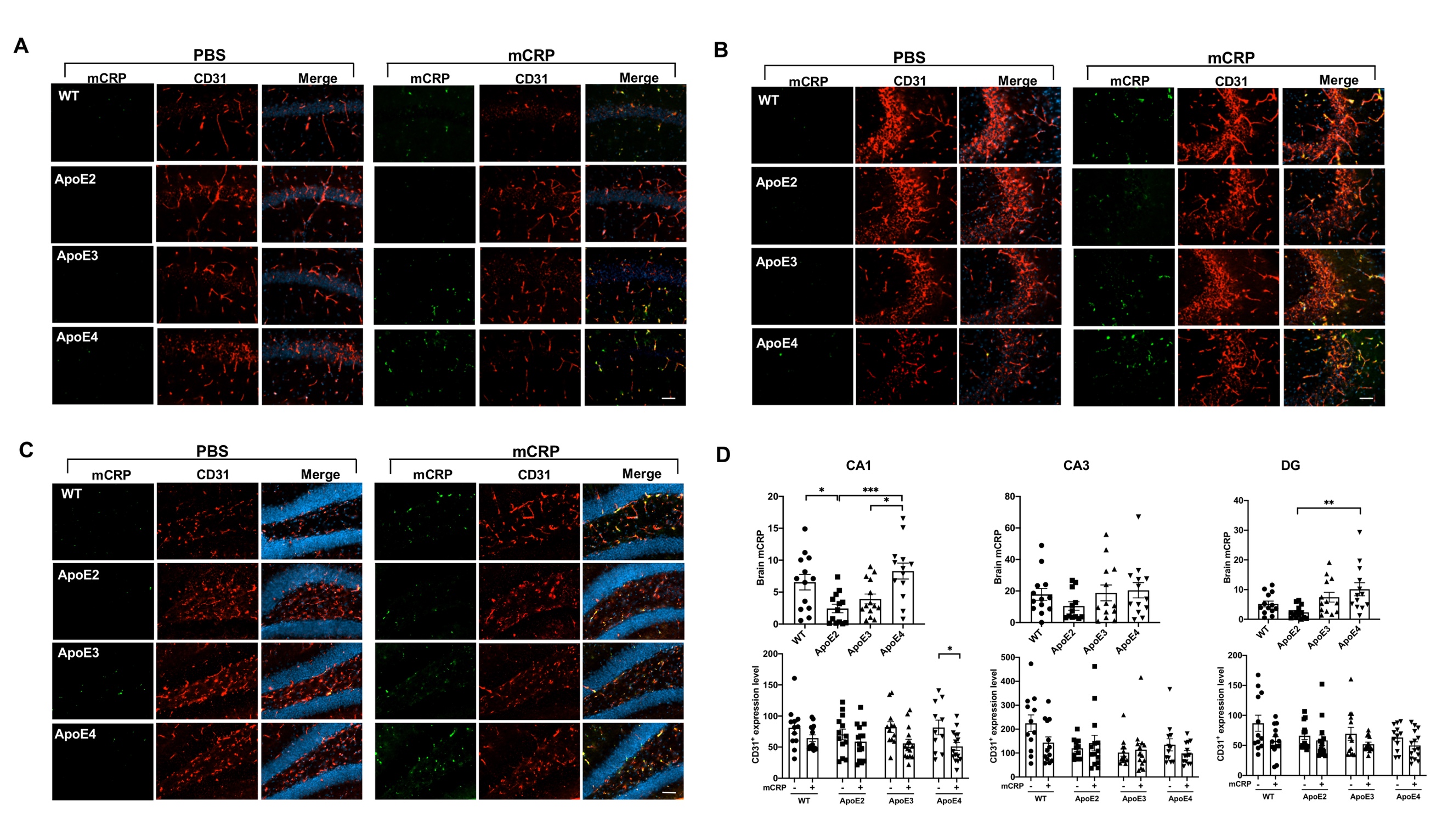

**Supplemental Figure 4**

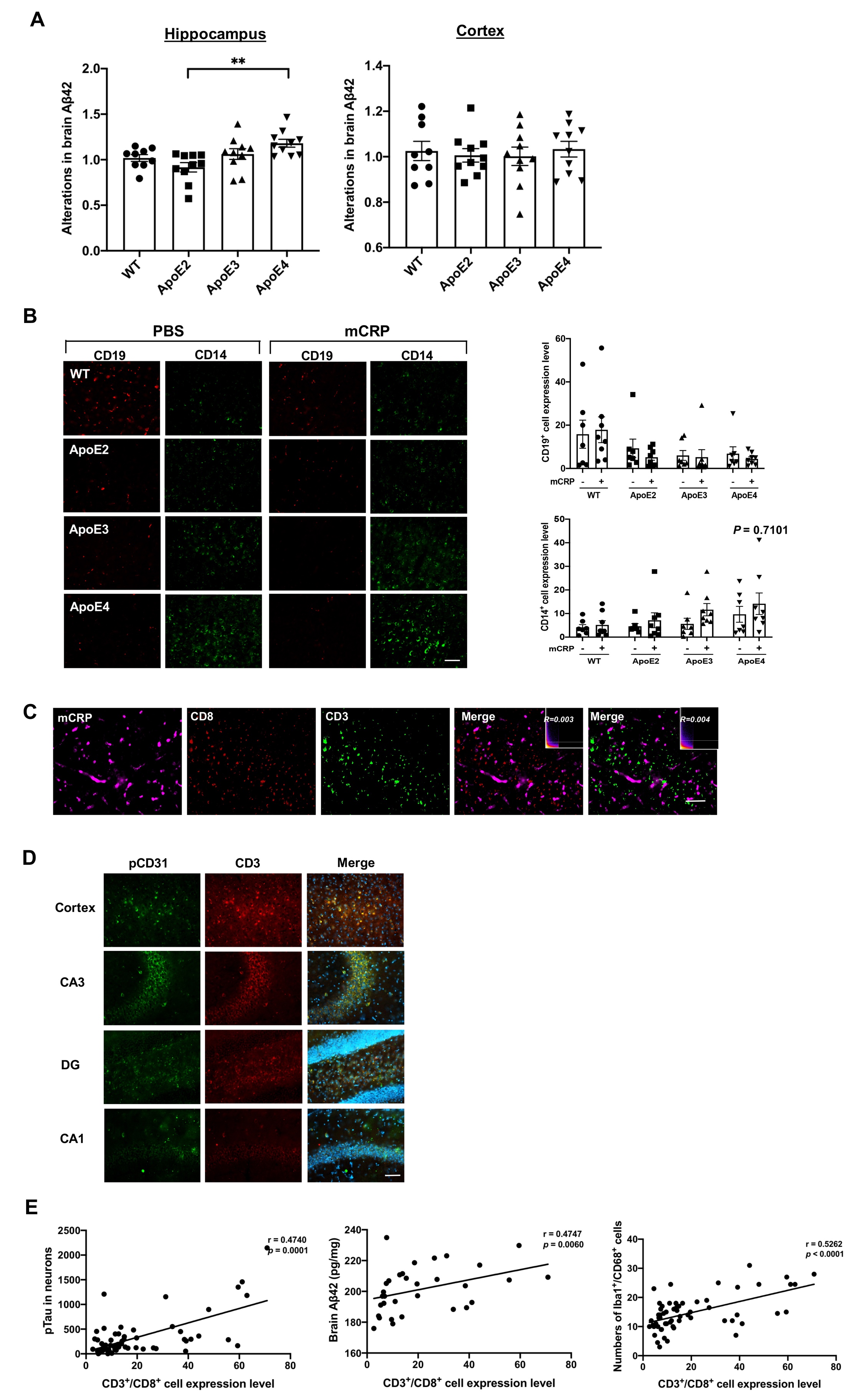

**Supplemental Figure 5**

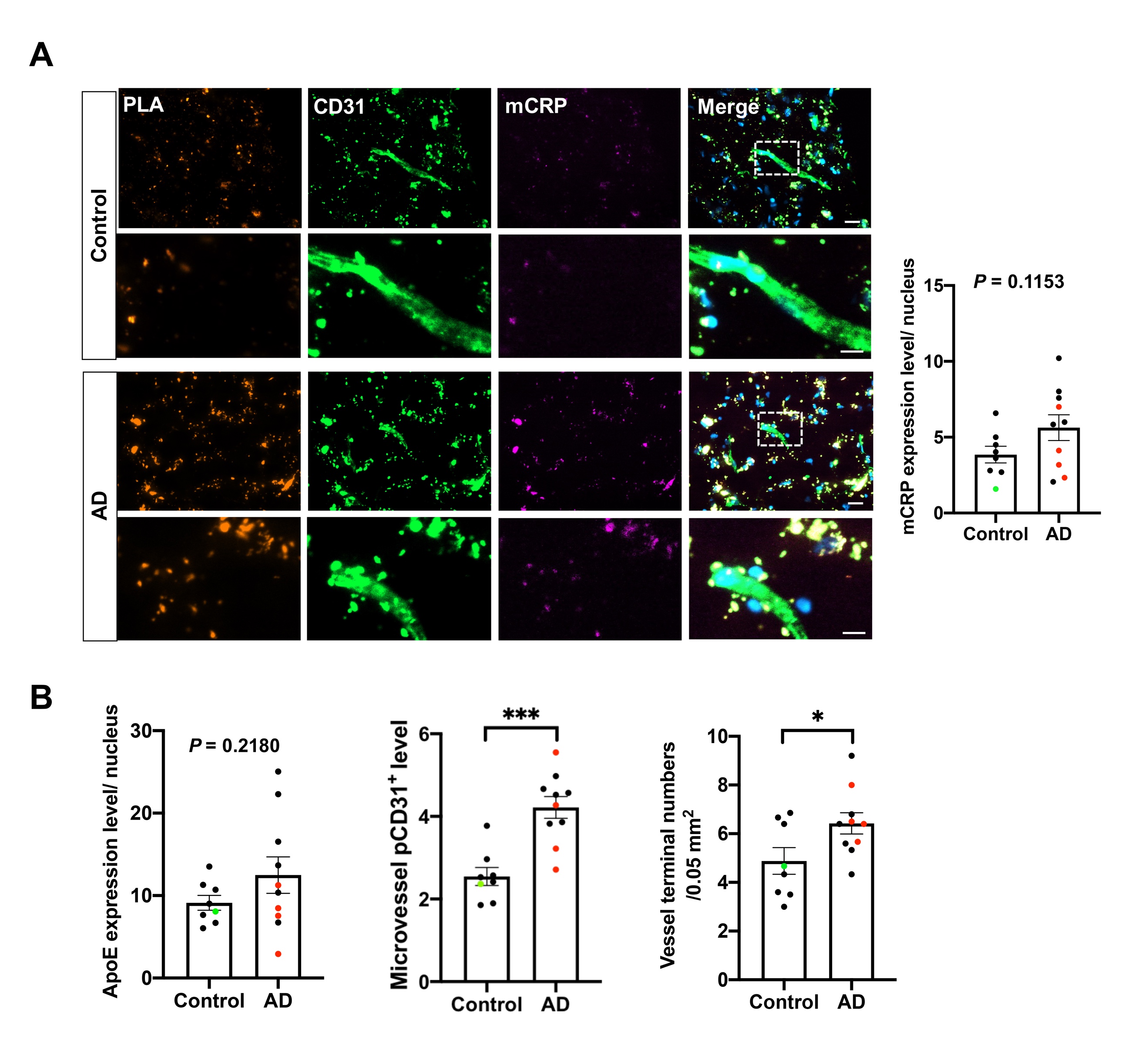

**Supplemental Figure 6**

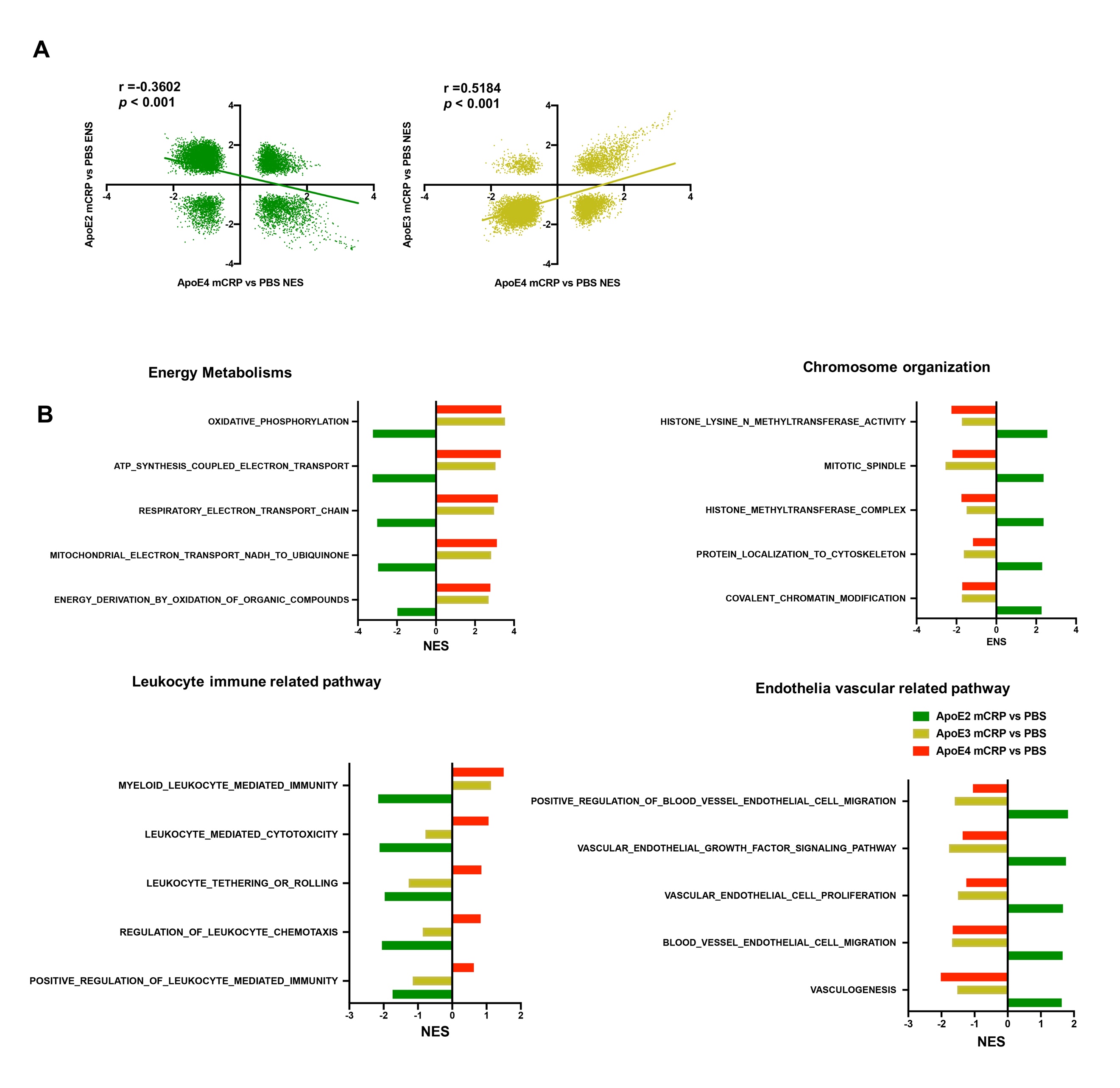
